# Climate change effects on terpene expression differ among populations and have consequences for herbivory in an invasive plant

**DOI:** 10.64898/2026.10.13.758875

**Authors:** Emma L. Boehm, Ryan Briscoe Runquist, David A. Moeller

## Abstract

1. Plant secondary metabolites often moderate interactions with abiotic and biotic factors. For example, secondary metabolites often deter insect herbivores from and influence the extent of damage to plant tissues. However, little is known about how climate change may alter the expression of secondary metabolites and the nature of the resulting effects on plant-herbivore interactions. As climate change continues to shift environmental conditions away from historical norms, climate change effects on plant-herbivore interactions may have important consequences for the range expansion of invasive plants and the efficacy of biocontrol agents across the range.
2. In this study, we examined climate change effects on terpene expression in common tansy (*Tanacetum vulgare*) in four field transplant sites in Minnesota, U.S.A. that include experimental manipulation of precipitation and temperature. Each transplant site included genotypes from 10 natural populations that span climate variation across Minnesota. We quantified leaf terpene profiles in two years (2024 and 2025) using targeted gas chromatography-mass spectrometry. We surveyed insect herbivory to the same genotypes in experimental field sites to ask how terpene profiles and chemotypes contribute to the likelihood of herbivory.
3. We detected evidence that climate change manipulation altered terpene production and depended on source population. Similarly, the extent of temporal variation in terpene profiles depended upon source population. We also found variation in terpene profiles among sites, further emphasizing the role of the environment in shaping terpene production. These shifts in terpene profiles, in turn, influenced the likelihood of herbivory. Herbivory was higher at the southern compared to northern sites and susceptibility to herbivory was higher in the northern compared to southern populations, particularly at northern sites where average herbivory was lower. The likelihood of herbivory also varied among chemotypes and the frequency of chemotypes varied among source populations, suggesting a complex interplay between the historical and current environments, terpene production, and plant-herbivore interactions.
4. **Synthesis.** Together, our results suggest a combined role of the abiotic environment and population differentiation in moderating plant-herbivore interactions via shifts and variation in terpene profiles, which may influence invasive range dynamics and complicate the regional management strategies of common tansy.

## INTRODUCTION

Invasive species often encounter novel abiotic and biotic environments in their introduced ranges (Mooney and Cleland 2001, Prentis et al. 2008). Considerable research has been dedicated to examining ecological and evolutionary responses to novel environments and the consequences for invasion dynamics (Moran and Alexander 2014, Colautti et al. 2017). In plants, responses to abiotic and biotic stress often involve the production of phytochemical compounds, especially plant secondary metabolites (Moore et al. 2016). Secondary metabolites have most commonly been shown to deter insect herbivores (reviewed in Afroz et al. 2021), but they may also modulate responses to abiotic factors (e.g. drought and cold stress: Llusià and Peñuelas 1998, Zhao et al. 2020, Jogawat et al. 2021), which may be particularly important as climate change causes more extreme and unpredictable environments (Gallagher et al. 2010, Akbar et al. 2024). For example, increased production and accumulation of secondary metabolites (specifically flavonoids) enhanced radical scavenging and prevented water loss in *Arabidopsis thaliana* (Nakabayashi et al. 2014). Given that secondary metabolite expression may be context-dependent in relation to the abiotic environment, it may be expected that secondary metabolite production between populations reflects climatic differences along latitudinal gradients in the geographic range (Sampiao et al. 2016, Liu et al. 2021, Xiao et al. 2020). Additionally, abiotic differences across the geographic range may shape biotic interactions and geographic variation in herbivore pressure may alter secondary metabolite production (Cao et al. 2025), forming a complex dynamic between abiotic and biotic factors and secondary metabolites.

Terpenes and their modified forms (i.e. terpenoids) are the largest class of plant secondary metabolites involved in both physiological processes and the mediation of abiotic and biotic stressors (Chatterjee and Pal 2023). In many plants, terpenes and terpenoids (hereafter, collectively referred to as “terpenes”) confer defense against herbivores (Levin 1976, Moore et al. 2016) and/or attract insect predators (Mumm et al. 2008, Unsicker et al. 2009). Terpenes have also been shown to increase plant performance under temperature or water stress in controlled environments (Loreto et al. 1998, Zhao et al. 2020; reviewed in Chatterjee and Pal 2023, He et al. 2023). Furthermore, because chemical pathways for responses to abiotic and biotic stressors are often shared, the production of certain terpenes may provide direct and indirect mitigation of a wide variety of environmental stressors (Pichersky and Raguso 2018). However, secondary metabolite production may be costly resulting in energetic tradeoffs between the production of terpenes and growth and reproduction, particularly in “stressful” environments (Züst and Agrawal 2017; Redman et al. 2001, Neilson et al. 2013). For example, in conifers, the emission of terpenes related to abiotic stress may reduce investment in defense compounds and increase susceptibility to insect herbivores (Holopainen et al. 2013). However, the expression of terpenes in relation to (manipulated) abiotic variation in natural field conditions remains poorly understood (reviewed in Bourtsoukidis et al. 2025; but see Pratt el al. 2014, Sun et al. 2022, Dobhal et al. 2024, Nomoto et al. 2025). In particular, the effects of climate change on terpene expression in the field have rarely been studied (reviewed in Jamieson et al. 2017; but see Gargallo-Garriga et al. 2017, Kivimäenpää et al. 2022).

It can be inferred that climate change may have complex effects on terpene production based on responses to individual abiotic stressors (Yuan et al. 2009). Elevated temperatures may increase terpene synthesis by promoting enzymatic activity (Liu et al. 2021, Jamloki et al. 2021) whereas reduced precipitation may lower terpene synthesis by restricting stomatal conductance and photosynthesis (Llusià and Peñuelas 1998, Kreuzwieser et al. 2021). Though still an emerging area of research, global change factors likely also interact in synergistic or antagonistic ways to shape production of secondary metabolites (Jamieson et al. 2017). Relatedly, there may be upper limits on the production of terpenes in highly stressful environments (Llusià and Peñuelas 1998, Holopainen et al. 2013). However, metabolomic responses to climate change factors are often complex and the consequences of altered terpene expression for plant-herbivore interactions remain unclear (reviewed in Jamieson et al. 2017 and Sun and Fernie 2024).

Common tansy (*Tanacetum vulgare*) is a plant native to Eurasia that was introduced to North America (Clasen et al. 2011). In the native range, terpene profiles vary geographically, deter insect herbivores, and attract predators of insect herbivores (Keskitalo et al. 2001, Benedek et al. 2015, Clancy et al. 2016, Rahimova et al. 2024). Chemical profiles are often dominated by one to three prominent compounds, comprising 40-99% of the profile, forming ecologically relevant “chemotypes” (Müller et al. 2026). There is also some evidence that increased production of certain terpenes may reduce growth and delay flowering time in common tansy, suggesting there may be tradeoffs associated with producing specific terpenes or terpene profiles (Keskitalo et al. 2001). Furthermore, chemotypic variation among native populations is sometimes associated with geographic variation in abiotic environments (Müller et al. 2026). Although most chemical profiling in common tansy has been conducted in the native range, invasive populations have been shown to produce more terpenes than native populations (Wolf et al. 2011). However, studies have not yet considered the ecological consequences of variation in terpene expression for invasion dynamics in different climate scenarios.

To determine how climate change alters the expression of terpene profiles and plant-herbivore interactions in invasive common tansy, we conducted a transplant experiment with manipulations that simulated climate change. Four transplant common gardens spanned two key climatic gradients in a region of the United States with a high density of invasive populations and observations of recent range expansion (state of Minnesota: temperature declines south to north, and precipitation declines east to west). Ten source populations from across Minnesota were planted into each common garden, where we also manipulated temperature and precipitation to simulate projected climate change. After three growing seasons, we quantified leaf terpene profiles for all ten populations in climate change (elevated temperature and reduced precipitation) and ambient (control temperature and control precipitation) treatments. We were particularly interested in testing the hypothesis that evolution during invasion has caused populations to respond differently to climate change (i.e. population × treatment interaction for terpene expression). We also quantified terpenes over two years to evaluate the extent to which environmental variation over time modulates terpene production. Last, we quantified leaf herbivory by insects in two years to ask whether herbivory varied geographically across the region and was related to variation in terpene expression (chemotypes).

## MATERIALS AND METHODS

### Field transplant experiment and climate change manipulations

We collected seeds in fall 2019 from ten common tansy populations across four regions (northwest = NW, northeast = NE, southwest = SW, southeast = SE) in Minnesota, spanning strong gradients in temperature and precipitation (Table S1). We grew 5-20 maternal families per population (depending on population size) in the greenhouse to standardize maternal environmental effects (Comstock and Robinson 1948). We randomly crossed among genotypes within populations to generate seeds for the field experiment and stored seeds at 4.

In 2022, we germinated seeds on a greenhouse mist bench for one week, transplanted seedlings into 2” pots, and grew them in a greenhouse until most plants had one true leaf. In spring 2023, plants were placed in an uncovered outdoor cold frame before transplanting into field sites. When most plants had 2-4 true leaves, we transplanted seedlings into the rectangular mesh pots (25.4 x 25.4 x 15.24 cm; Total Pond, Aurora, OH) lined with landscape fabric that were already installed at each common garden and filled with field soil. These measures mitigated unintended spread of common tansy in the field. We watered seedlings for two weeks and replaced individuals that died within the first two weeks.

The four common gardens were established in four distinct combinations of temperature and precipitation in Minnesota (NW, NE, SE, SW; Fig. 1). We also implemented fully factorial manipulations that simulate climate change conditions for both temperature and precipitation. To simulate elevated temperatures, we used passive open-topped warming containers (OTCs) constructed using four clear polycarbonate panels (Tuftex PolyCarb, Kutztown, PA). The warming effect of the OTC design was validated in a common garden experiment (see

**Fig. 1.**
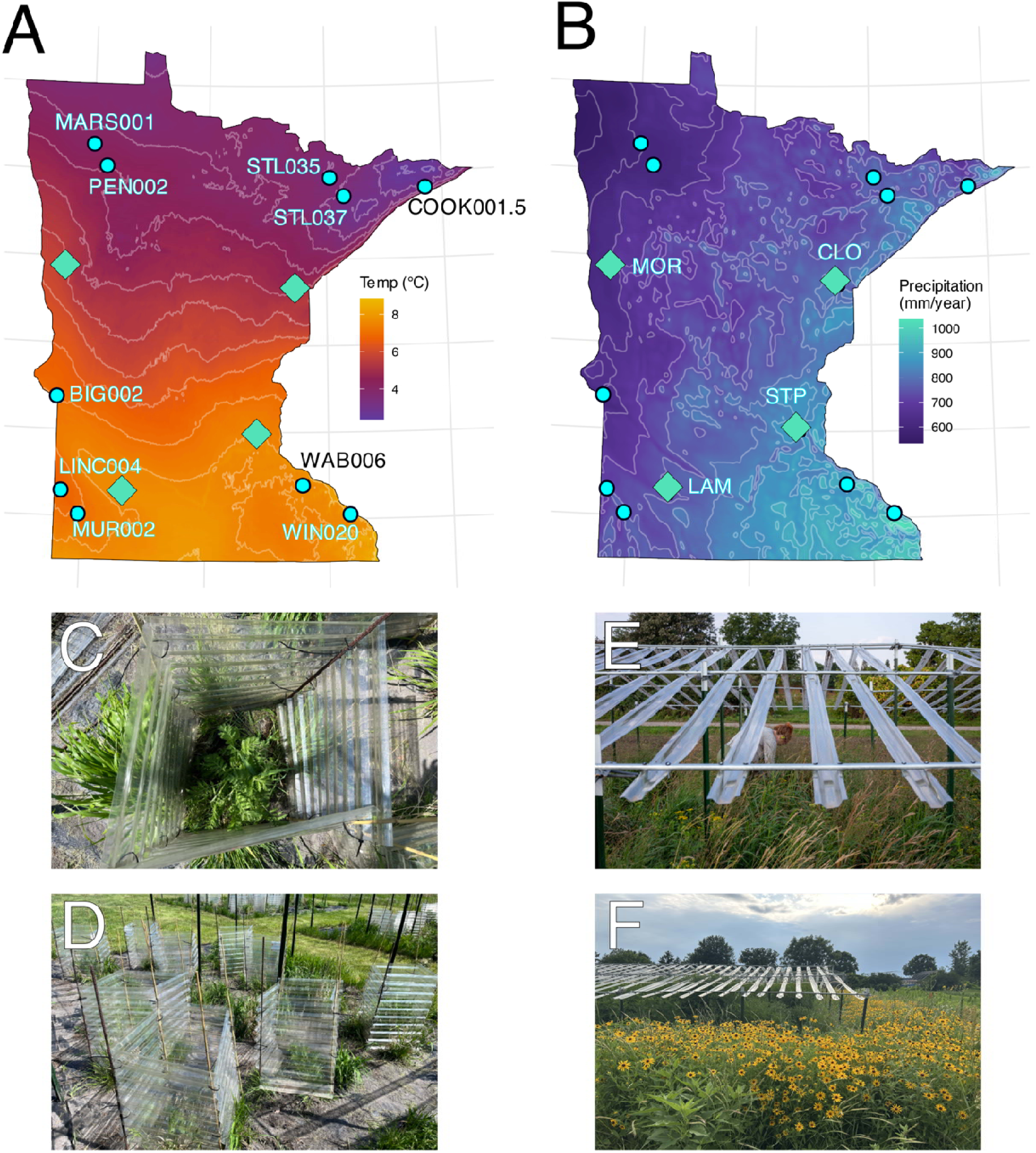
Focal populations, transplant sites, and experimental design with climate change manipulations. (A) Mean annual temperature of focal populations (circle) and transplant sites (diamonds). (B) Cumulative annual precipitation of focal populations (circle) and transplant sites (diamonds). (C-D) Manipulations that simulate projected elevated temperature in Minnesota. (E-F) Precipitation reduction treatments that simulate predicted precipitation changes in Minnesota.

Briscoe Runquist et al. 2026). The OTCs increased temperature by 2-4°C (day) and 0-1°C (night), closely simulating climate change predictions for the region (Liess et al. 2022, Ford et al. 2021, Chen and Ford 2023). To simulate the predicted decrease in average regional precipitation (Arias et al. 2021), we used rainout shelters where ca. 35% of precipitation was removed by clear plastic gutters (modified from Rudgers et al. 2023). In ambient precipitation plots (control), gutters were upside down such that precipitation was largely not removed from plots. Shelters were arranged in a split-plot design where one side of each plot received ambient precipitation and the other simulated precipitation reduction (5 replicate blocks per transplant site). We quantified terpene profiles and herbivory for individuals in ambient plots (control temperature and control precipitation) and those with both climate change manipulations (elevated temperature and reduced precipitation) (N = 10 source populations × 4 transplant sites × 2 treatments × 2-4 individuals per source population per treatment per transplant site = 310).

### Metabolomics

We collected leaf tissue from individuals in two years. In 2024, we collected samples from all control-control individuals in our two northern sites (NW and NE). In 2025, we collected samples from all four common gardens and for the elevated temperature-reduced precipitation treatment combination and the ambient treatment. We stored leaf tissue at -20 until terpene extraction.

For terpene extraction, we flash froze tissue using liquid nitrogen and ground samples using flash frozen ceramic mortars and pestles (CryoGrinder, OPS Diagnostics, Lebanon, NJ, USA). We transferred ground tissue (50 mg) to amber autosampler vials (2 mL, Agilent Technologies, Inc., Santa Clara, CA, USA) and added hexane that contained a standard (1 mL of 2 µg/mL Borneol-D3 in 2024, 10 µg / mL Limonene-D5 in 2025). Samples were stored at 4 for 24 hours, supernatant was transferred into clean vials, and extractions were stored at -80. The University of Minnesota Center for Metabolomics and Proteomics performed Gas Chromatography-Mass Spectrometry (GC-MS), peak detection and integration, data preprocessing, and metabolite identification. Peak sizes were normalized by dividing raw peak area by the internal standard peak area. We only considered a compound signal as present if it was detected at an intensity of at least ten times the blank background control measurement. We also ran three samples in triplicate and found all normalized peak areas for each metabolite within triplicate samples were highly correlated (all r > 0.99, p < 0.001).

### Quantifying herbivory in the field

We quantified leaf herbivory for all individuals in 2025. We surveyed leaf damage on four branches per plant at each common garden (mean leaves surveyed per plant based on subset = 148). Common tansy may be damaged by insects with diverse feeding habits so we assessed leaves for sucking, chewing, or and boring damage. We scored plants as having herbivory if ≥ 4 leaves had damage. We excluded four individuals from analyses with evidence of substantial mammalian herbivory (e.g., stem removal).

### Statistical analyses

All analyses were performed in R (version 4.3.1) and RStudio (version 2023.12.1+402). We used ggplot2 to produce graphs (Wickham 2009) and cowplot to create multi-panel figures (Wilke 2024). We used dplyr for data manipulation (Wickham et al. 2023). For linear models, we used ANOVAs with Type II sums of squares in the car package.

#### Environmental and genetic effects on terpene profiles

We first assigned compounds to compound classes (e.g. monoterpenoids) using the chemodiv package (Petrén et al. 2023). To account for heteroscedasticity and right-skewedness (a common outcome of the peak normalization procedure; De Livera et al. 2012), we log transformed data before analysis (Van den Berg et al. 2006, Kohl et al. 2012, Sun and Xia 2024). For methods involving Euclidean distance, we subsequently mean-centered and unit variance-scaled (mean = 0, standard deviation = 1) each metabolite (Anwardeen et al. 2023, Sun and Xia 2024). When metabolites were absent from an individual sample but present in the full dataset, we converted zeroes to an arbitrary small value (10), permitting the inclusion of all compounds in analyses (Sun and Xia 2024). Because autoscaling can reduce some biologically relevant differences in metabolite abundance, we also ran supplementary analyses of Pareto-scaled data (Sun and Xia 2024), which less severely reduces the effects of highly abundant compounds (results were qualitatively similar, see Supplemental Information).

To determine how population and environment (site and treatment) shaped terpene profiles, we evaluated differential terpene expression. We used a PERMANOVA with Euclidean distance (for quantitative expression data) and 999 permutations using the vegan package (Oksanen et al. 2025). We evaluated significance using type II tests with the RVAideMemoire wrapper package (Herve 2025). We included the fixed effects of source population, transplant site, treatment (control-control or elevated temperature-reduced precipitation), block nested within transplant site, and all two- and three-way interactions. We used the *pairwise.adonis* function in the pairwiseAdonis package (Martinez 2017) to perform post hoc pairwise PERMANOVAs with a correction for multiple comparisons using the Benjamini-Hochberg method. Following PERMANOVAs, we evaluated dispersion of individuals around the group centroids as a validation metric using vegan (*betadisper* function).

We used the *ancombc2* function in ANCOMBC (Lin and Peddada 2020) to identify differentially expressed compounds and applied additional specifications (removal of rare species and Benjamini-Hochberg correction) to account for pseudocounts that may inflate false discovery rates and abundance differences. We also evaluated differences in composition (i.e., compound presence/absence and relative frequency), but focus on terpene profiles in the main text (compositional results were similar, see Supplemental Information).

#### Temporal variation in terpene profiles

We evaluated temporal variation in terpene expression using data from the ambient treatment (control-control) from the NW and NE transplant sites for both 2024 and 2025. We removed novel terpenes discovered in 2025 (n = 7) and ran a PERMANOVA with source population, transplant site, year, population × site, population × year, and site × year as fixed effects. We used a PERMANOVAs with Euclidean distance (999 permutations) and evaluated significance with type II tests. As above, we present compositional differences in the Supplemental Information.

#### Determination of chemotypes

We classified individuals as chemotypes using hierarchical clustering based on Euclidean distance of all identified terpenes using the factoextra package (Kassambara and Mundt 2020). We used the elbow method to determine the number of clusters. We further validated the number of clusters using the NbClust package (Charrad et al. 2014), which summarizes k-cluster recommendations from multiple distance measures and clustering methods.

To determine which terpenes had the largest relative contributions to each chemotype, we performed a partial least square discriminant analysis (PLS-DA) followed by extracting Variable Importance in Projection (VIP) scores > 1 as well as the weighted sum of partial-least squares regression coefficients using MetaboAnalyst (version 6.0; see Xia et al. 2009). We also extracted the top three Z-scores directly from the hierarchical clusters to visualize differences in relative abundance, where higher Z-scores indicate increased compound expression (top three Z-scores > 0.5 for all chemotypes except for Chemotype 3).

We evaluated the fixed effects of site, population, treatment, and all two-way interactions on chemotypic variation using a multinomial logistic regression model specified using the *multinom* function in the nnet package (Venables and Ripley 2002).

#### Relationship between herbivory, climate change, and chemotypic variation

To evaluate how climate change and chemotypes influenced the likelihood of herbivory (binary metric), we used generalized linear models with a binomial distribution (*glm* function in base R). We included chemotype, source population, transplant site, treatment and two-way interactions as fixed effects. We excluded the chemotype × source population interaction because combinations of the two factors were highly unbalanced. We also examined variable inflation factors (VIFs) of main effects, which indicated that multicollinearity was minimal (all VIF < 2.7).

If we detected a significant main effect or interaction with chemotype, we repeated the PERMANOVA and ANCOMBC analyses separately for each chemotype to determine which terpenes differed in expression between predictor categories. We further tested whether herbivory varied with the latitude and longitude of source populations. We used a generalized linear model with population mean herbivory as a dependent variable and chemotype, site, treatment, latitude, longitude, and all two-way interactions (except for chemotype × source population) as fixed effects. We centered latitude and longitude (mean = 0) to facilitate comparison of regression coefficients.

## RESULTS

### Site, population, and treatment effects on terpene expression

In 2025, we characterized terpene variation across all four transplant sites and in the dual climate change and dual ambient treatments. We identified 51 compounds, which were comprised primarily of monoterpenes (49.0%) and sesquiterpenes (35.3%) (Table S2 for full details).

Terpene expression (Euclidean) was significantly different among sites (PseudoF_3,228_ = 7.99, p < 0.001; Fig. 2B, Table S3). More specifically, all site combination pairs differed from one another (adjusted p-values < 0.05) except for NE-SW. While population and treatment did not individually modulate terpene expression, we found evidence that climate change effects on expression depended upon source population (type of G x E interaction) (PseudoF_9,228_ = 1.50, p = 0.050; Table S3). Many, but not all, population by treatment combinations differed in terpene profiles (Table S4). Within these population-treatment pairs, we found more differences between regional groups (i.e. NW, NE, SW, SE) than populations within regional groups, further supporting a spatial pattern in terpene profile differentiation (Fig. 2C, Table S5). We did not detect a clear pattern of climate change effects within populations or regions (Fig. 2C). Dispersion of terpene profiles did not differ among sites, populations, or treatments.

**Fig. 2.**
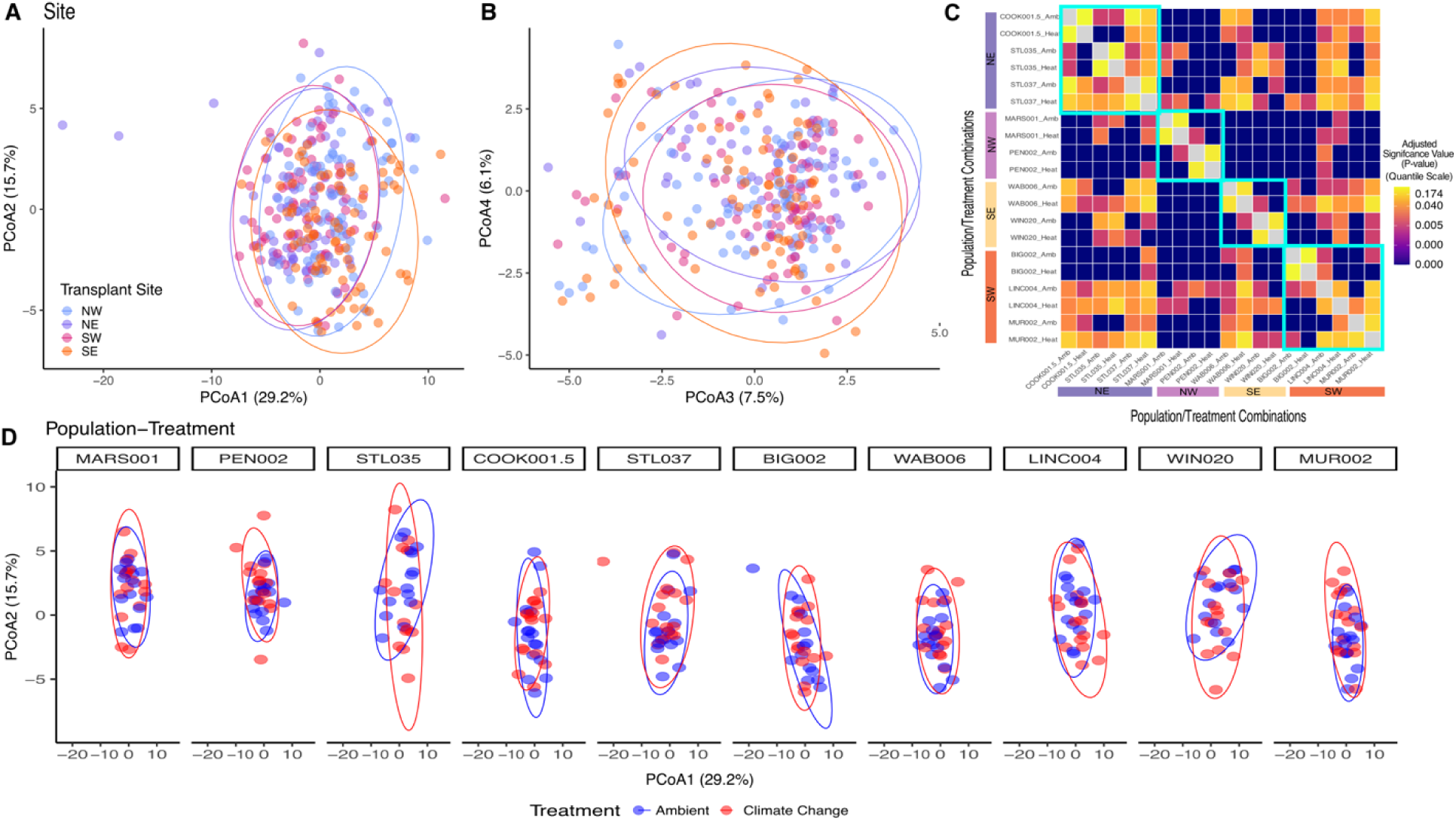
Visualization of (A-B) terpene profiles between transplant sites as well as (C-D) population-treatment combinations. Ellipses represent 95% confidence intervals. Colors are transplant sites in (A-B), adjusted p-value quantiles (C), and populations from north to south in (D). Note that the scale of the axes differs between panels.

The strongest evidence for differential expression of individual terpenes occurred between treatments. The climate change treatment substantially decreased the relative abundance of two compounds compared to the ambient treatment: cis-sabinol and isogermacrene-D (lfc = - 0.70 to -0.84, p < 0.001, all p-adjusted < 0.001). We detected five additional compounds with reduced expression in the climate change treatment and only one with increased expression; however, p-values were significant before but not after adjustment for multiple tests (all p < 0.05, all p-adjusted > 0.1; Fig. S1). While there was some suggestion of differential expression of 7 individual terpenes among sites and 6 among populations, p-values were significant before but not after adjustment for multiple tests (for more detail, see Fig. S2).

### Site, population, and treatment effects on chemotypic variation

In 2025, the elbow method and the ensemble approach (NbClust) both identified five chemotypes (Fig. 3A). The top two compounds differed for all chemotypes (Fig. 3B, Table S6, Table S7). For example, dominant compounds for Chemotype 4 were 3-car-en-5-one and cis-chrysanthenol-acetate; whereas, those for Chemotype 3 were alpha-thujone and beta-thujone (Table S6 & S7).

**Fig. 3.**
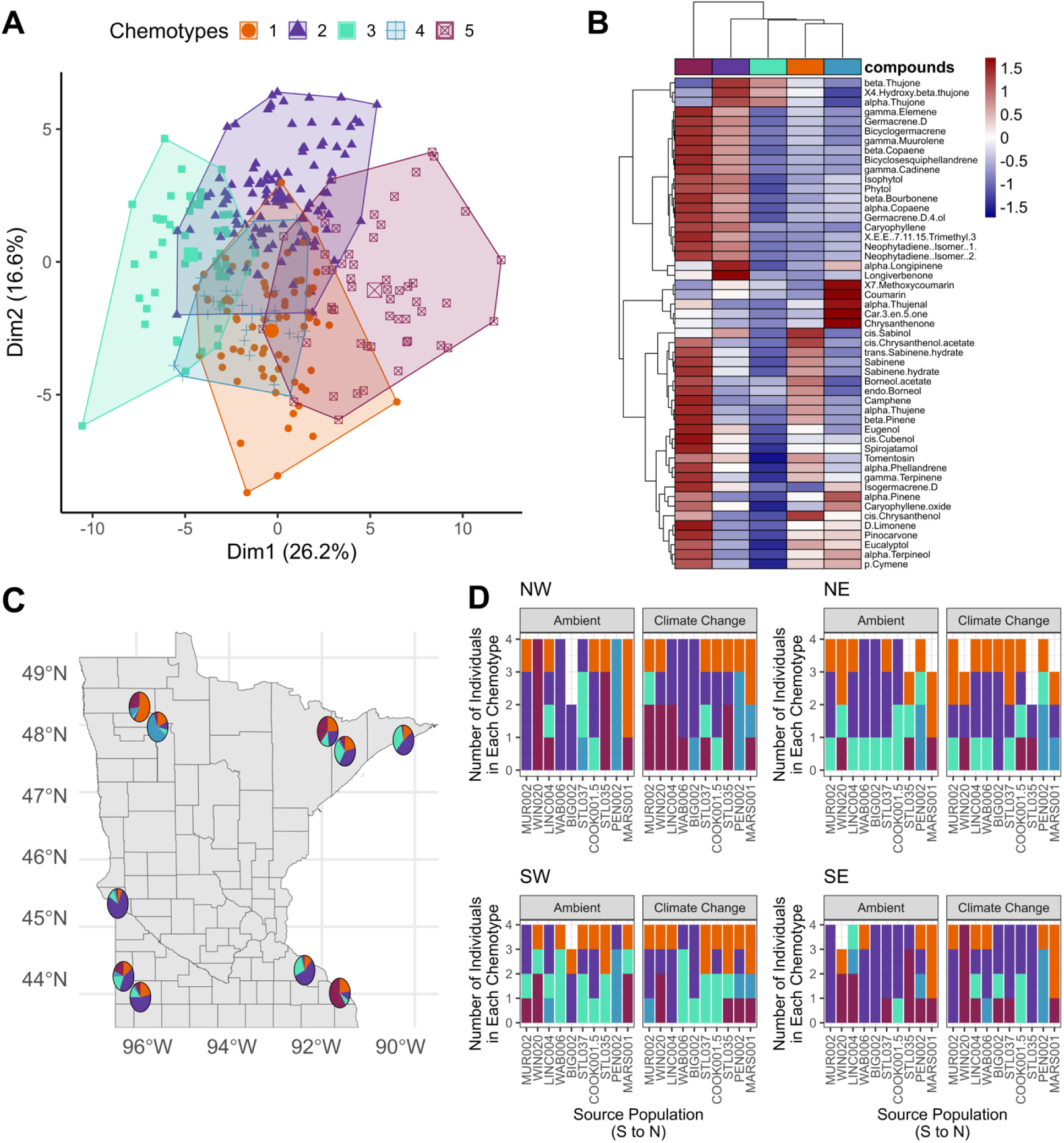
Cluster analysis of terpene expression revealed five distinct chemotypes composed of unique combinations of dominant and minor compounds. **(A-B)** that are distributed over source populations (C-D). Points in (A) and counts in (D) are all individuals characterized from 10 populations across four transplant sites. The gradient bar in (B) shows differences in relative abundance of terpenes (red = increased, blue = decreased). Colors are chemotype groups as determined by hierarchical clustering.

Chemotype composition differed among sites (LR ^2^ = 8.16, p = 0.0056; Fig. 3D, Table S8) and populations (LR ^2^ = 54.7, p < 0.001; Fig. 3C, Table S8). Specifically, the NE and SW sites had relatively more Chemotype 3 but fewer Chemotype 5 individuals compared to the NW and SE sites (Chemotype 3: 16-21 vs. 6-7 vs.; Chemotype 5: 6-10 vs. 18-20 individuals). For populations, chemotype composition differed for nine of 45 pairwise comparisons (all p < 0.05, Fig. S3). Perhaps unsurprisingly, the results for chemotypes mirror those for terpene composition in that composition was similar between NE and SW and between NW and SE but differed between the pairs (see Supplemental Information for more details).

### Temporal variation in terpene expression

We identified 44 core compounds that were present in both 2024 and 2025 (56.8% monoterpenes, 31.8% sesquiterpenes, 4.5% diterpenes, 4.5% coumarins, 2.3% phenylpropanoids). Seven additional compounds were identified only in 2025: these novel compounds included sesquiterpenes (4), diterpenes (2), and one hydrocarbon.

In an analysis that combined both years of data for the NW and NE sites, we found that differential expression in the two years depended upon population and differed among sites (population × year interaction: PseudoF_9,137_ 1.45, p < 0.01, site: PseudoF_1,144_ = 6.05, p < 0.001; Table S9). Chemotypes within populations were more labile between years for certain populations: we found that some populations had an increased likelihood of switching chemotypes between years (LINC004) whereas others (MAR001) remained relatively static (Fig. S4). The most common chemotype change between years was from Chemotype A to Chemotype C or D, both of which tended to produce fewer, targeted compounds (Fig. S4).

For individual terpenes, we identified 17 terpenes that were differentially expressed between years. There was no apparent overall tendency for expression to be increased vs. decreased between years: nine compounds had higher expression (alpha-longipinene, alpha-thujenal, alpha-thujone, borneol-acetate, car-3-en-5-one, chrysanthenone, cis-chrysanthenol-acetate, longiverbenone, spirojatamol) and eight compounds had lower expression (alpha-phellandrene, alpha-pinene, beta-copaene, bicyclosesquiphellandrene, eugenol, gamma-muurolene, sabinene, sabinene-hydrate) in 2024 compared to 2025 (all p < 0.01, all p-adjusted < 0.05). Of the 100 pairwise comparisons of populations, 70 differed between years but were not significant after p-value adjustments (all p < 0.05, p-adjusted > 0.1).

There were five compounds (alpha-copaene, beta-copaene, bicyclosesquiphellandrene, gamma-cadinene, and germacrene-D) that had increased expression at the NW site compared to the NE site (lfc =0.53 to 0.086, all p < 0.01, all p-adjusted < 0.05).

In the combined dataset, the elbow method identified four chemotype clusters years (supported by NbClust; Fig. S5A, Fig. S5B). The proportion of individuals in each chemotype differed between years and sites (Fig. S5C).

### Relationship between herbivory, climate change, and chemotypic variation

There was a higher likelihood of herbivory at southern transplant sites compared to the northern transplant sites (χ^2^ = 58.4, p < 0.001; Fig. 4, Table S9). The effect of the climate change treatment on herbivory differed among chemotypes (chemotype × treatment: χ^2^= 11.5, p < 0.05; Fig. 4, Table S10). In particular, two chemotypes (3 and 4) had an increased likelihood of herbivory under climate change while the other chemotypes exhibited little or no differences between treatments. Climate change was also associated with higher mean herbivory at the NE site where the overall likelihood of herbivory was lowest (37%) compared to other sites where herbivory was higher for all chemotypes (58-93%; transplant site × treatment: χ^2^ = = 11.8, p < 0.01). Importantly, the effect of climate change on herbivory depended on population (increased damage for three of ten populations: population × treatment: χ^2^ = 23.4, p < 0.01; Table S10). Of the three populations, two were southern populations (MUR002, LINC004) and the other was a northern population (COOK001.5).

**Fig. 4.**
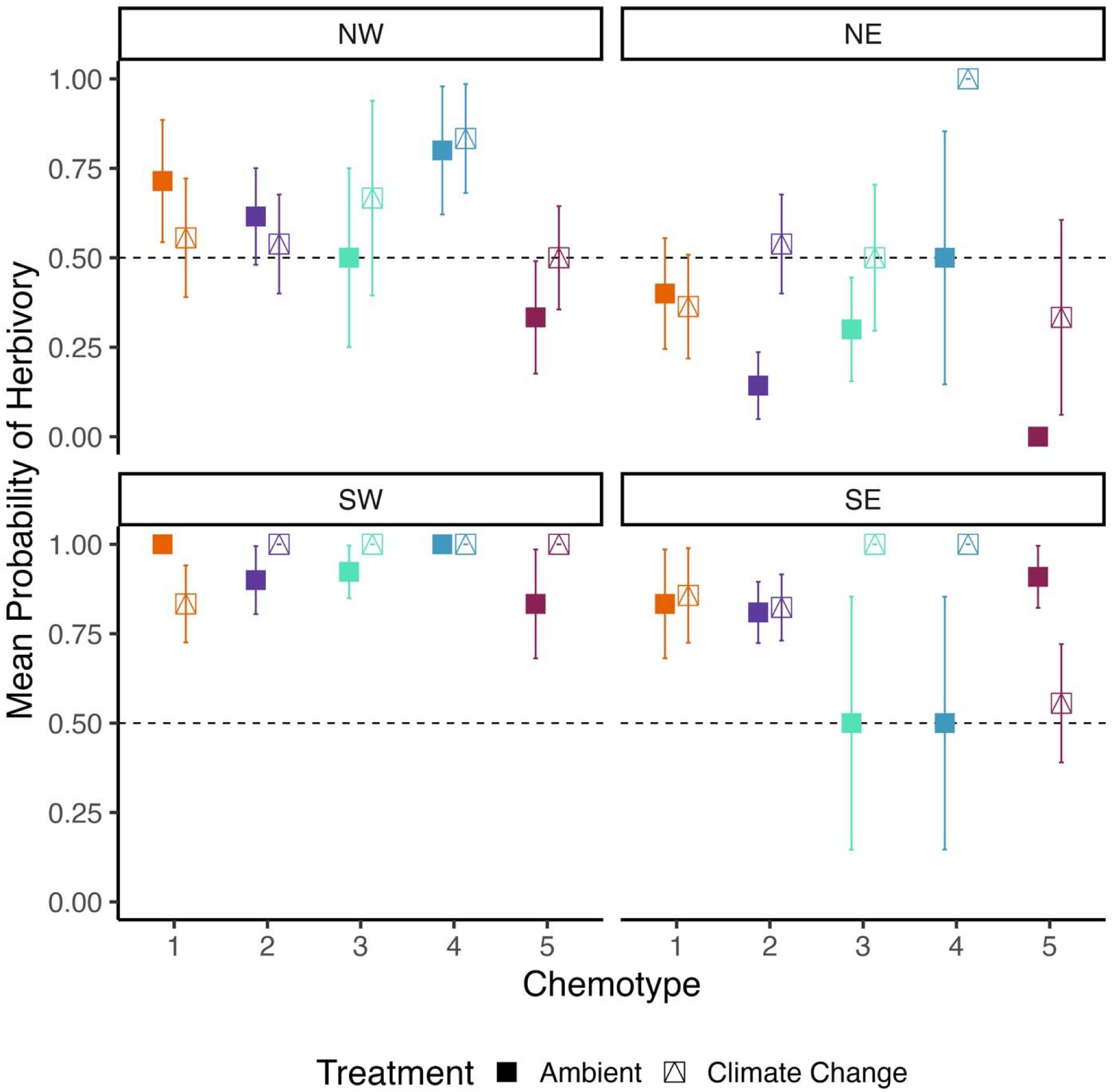
The effects of climate change on the likelihood of herbivory depended on chemotype. Points are means and bars are standard error. Transplant sites are denoted in panel labels (NW, NE, SW, SE). Colors are chemotype groups and shapes are field treatments (filled square: ambient treatment (control-control); open square: climate change = elevated temperature-reduced precipitation). Dashed line at 0.5 is for visual reference.

For individual terpenes of Chemotype 3, climate change reduced versus increased the expression of three and two compounds, respectively. Plants with Chemotype 3 expressed less beta-pinene, endo-borneol, and spirojatamol and more cis-sabinol and cis-chrysanthenol in the climate change treatment (all p-adjusted < 0.05). We did not detect differential expression of individual terpenes between treatments in Chemotype 4 after adjusting p-values.

Last, mean herbivory was greater for populations from northern compared to southern latitudes. Northern populations have greater susceptibility to herbivores compared to southern populations and the degree of latitudinal effects on the likelihood of herbivory depended on site (NW – SW, NE – SW, NE – SW, SW – SE pairwise contrasts: all p < 0.05; site × latitude: ^2^ = 7.99, p = 0.046; Fig. S6, Table S11). The pattern of greater resistance in southern populations is consistent with the finding that herbivore pressure was greatest at southern sites. We did not detect any interactions between latitude and chemotype or any significant effects of longitude (all p > 0.13; Table S11).

## DISCUSSION

Plant secondary metabolites often mitigate the effects of abiotic stressors and confer defense against herbivores and pathogens. At the same time, abiotic and biotic factors also influence the expression of secondary metabolites, which in turn may affect plant performance. Given that climate change is expected to increase abiotic stress in many environments, it may also have important consequences for plant-herbivore interactions via altered expression of secondary metabolites. In this study, we investigated the environmental and genetic contributions to variation in terpene expression using a transplant experiment coupled with manipulations that simulated climate change. In particular, we asked how climate change affected the profile of leaf terpenes and examined its consequences for herbivory in the invasive plant, common tansy. Environment strongly shaped terpene composition and profiles, both in terms of transplant site effects and differences between years. Importantly, we found that responses to climate change treatments depended upon source population, suggesting that climate change effects may not be predictable across the region despite the recency of invasion. Moreover, climate change increased the likelihood of herbivory for certain chemotypes while others exhibited little or no difference between treatments. Overall, our results suggest that climate change may have important consequences for plant-herbivore interactions by modulating the expression of terpenes.

### Environmental variation modulates expression of terpenes

The production of terpenes is inherently linked with the broader environment as environmental shifts often catalyze metabolomic changes (Byrd et al. 1999, Boncan et al. 2020, Keefover-Ring 2022, Felisberto et al. 2024). Our results similarly show that leaf terpene composition and profiles vary among transplant sites (differences in spatial environment) and years (differences in temporal environment). Together, the spatial and temporal variation in terpene production suggest that environmental filtering occurred between sites and years. Aside from representing different combinations of abiotic factors, transplant sites also serve as a proxy for potential climate change scenarios. Effects of temperature and precipitation on terpene production suggest that and that future climate variation will likely alter terpene profiles in common tansy.

Beyond average temperature and precipitation, the transplant sites or years may also differ in other abiotic and biotic factors. For example, the synthesis of many secondary metabolites may be influenced by soil type and nitrogen availability (Ormeño et al. 2008, Sanchez et al. 2008, Obata and Fernie 2012). In relation to other potential differences among sites, we also detected among-site differences in dispersion from the compositional analyses. This was primarily driven by increased compositional homogeneity of metabolites at the SE transplant site compared to the other sites. Variation in secondary metabolite production within (in addition to between) sites likely depends on environmental conditions as well (He et al. 2025), although the specific mechanisms of increased homogeneity at this particular site require further study.

Site variation in terpene expression did not simply reflect additive effects of temperature and precipitation on terpene production in common tansy. Though terpene profiles differed among transplant sites, the NE and SW sites were often more similar to one another than the other sites and shared similarities in both terpene profiles and chemotypes. The NE and SW sites are at opposite ends of historical climatic gradients, particularly in terms of average temperature, and are potentially representative of environments encountered in different stages of the invasion process (e.g., colonization vs. spread). Differential abiotic extremes may limit the diversity of terpene production and increase secondary metabolite profile similarity (Kaplan et al. 2004). However, it is more likely that individuals at the NE and SW sites experienced similar abiotic environmental “stress” within our experiment years that led to overlap in terpene production: the maximum temperatures and estimated evapotranspiration were similar at the NE and SW sites across the growing season (Fig. S7). Temperatures in the summer were particularly high for NE site in 2025 compared to historical averages (Fig. S8). Overall, these results suggest that terpene production is not only dependent on evolutionary responses to historical climatic variation but that weather in individual years may also lead to differential plasticity in terpene production.

### Climate effects on terpene expression vary among populations

To our knowledge, there are few other field studies that evaluate the effects of genotypic and environmental variation on terpene production (Pratt et al. 2014, Dobhal et al. 2024, Nomoto et al. 2025) and only one that also evaluates plant-herbivore interactions (Sun et al. 2022). Our study is unusual given that populations varied in their terpene profile responses to the climate change treatments. This suggests that evolutionary adaptation and potentially phenotypic plasticity (acclimation to environmental stress; Wada and Sewall 2014) across geographic clines may contribute to terpene production. The lack of a clear directional pattern of climate change effects within regions or populations further emphasizes the complexity of variation in terpene production. Environmental gradients have been shown to differentially shape both adaptation and plasticity of phytochemical responses to particular abiotic factors such as drought (Moore et al. 2014, Kessler and Kalske 2018, Gitonga et al. 2022, Castells and Sanchez Martinez 2025). Overall, the complexities of geographic and population differentiation in terpene profiles as well as differential responses to environmental effects may complicate management of this invasive species.

### Climate change effects on herbivory

It has been argued that biotic interactions are stronger at lower than higher latitudes and some past results support this hypothesis (reviewed in Schemske et al. 2009; but see Moles et al. 2011). As a result of the increased herbivore pressure at lower latitudes, more southern populations may evolve increased defense against herbivores compared to northern populations (Rasmann and Agrawal 2011). Our study is consistent with this hypothesis in that we found a higher likelihood of herbivory at southern sites and greater resistance to herbivory in southern source populations. These results are intriguing because while this pattern has been shown previously, tests of this hypothesis are often conducted on much larger, continental scales (Hargreaves et al. 2019; but see Kent et al. 2020). For example, Baskett and Schemske (2018) found that herbivory to *Phytolacca americana* was greater in southern North America and that southern populations were less palatable to herbivores in laboratory trials. Our study suggests that latitudinal gradients in biotic interactions are also important at finer spatial scales. In addition, our finding that chemotypic variation is related to herbivore damage in common tansy implicates terpenes as a potential target of selection in the southern part of the focal range of our study. Our results are surprising given that our study focused on an invasive plant with recent range expansion at southern range margins (Briscoe Runquist et al. 2026).

Despite these results, we found that populations consistently maintained a diverse array of chemotypes. More specifically, at least four of five chemotypes were present in all populations. The discrepancy between geographic variation in herbivory (and parallel population variation in resistance to herbivory) may be explained by several non-exclusive hypotheses.

First, differentially chemotypic variation may be maintained if there is conflicting selection on terpene expression by biotic and abiotic factors (Carley et al. 2021, Wittmann and Bräutigam 2025). For example, precipitation has been shown to influence phenotypic selection on terpene expression; selection favored a lower abundance of monoterpenes in wet conditions but increased abundance in drier conditions (Pratt et al. 2014). Similarly, allelic variation underlying glucosinolate (another group of secondary metabolites) production in *Boechera stricta* was maintained across populations due to pleiotropic effects on defense against herbivores and drought (Carley et al. 2021). Diversity of underlying compounds (and resulting chemotypes) also may be maintained as they collectively increase defense against single and simultaneous environmental factors (Jamieson et al. 2017, Whitehead et al. 2021). Regardless of the mechanism, high chemotype variation within populations has implications for management via biocontrol. In particular, if the efficacy of biocontrol agents (stem mining weevil, *Microplontus millefolii*, for common tansy; Gassmann et al. 2010) varies among chemotypes, then management may require a combination of approaches.

### Environmentally-driven shifts in individual terpenes

We found that individual terpenes varied in response to environmental variation: between treatments, among sites, and between years. In each of those cases, bicyclic monoterpenes and monoterpenoids (especially thujones, sabinenes, pinenes, carenes, and borneols) often exhibited shifts in expression. One reason that we may have detected plastic production of bicyclic monoterpenes in response to environmental variation is due to the rapid synthesis of these terpenes. Compounds of these families are synthesized using the pre-cursor geranyl diphospate (GPP) along the same methylerythritol phosphate pathway (MEP) (Degenhardt et al. 2009). Compared to heavier sesquiterpenes (with more core isoprene units), bicyclic monoterpenes are synthesized in fewer steps and require less carbon and metabolic energy to synthesize (Gershenzon 2017, Lanier et al. 2023). While some sesquiterpenes also varied between environments, these changes were more often between years (rather than sites or treatments) in our study. This aligns with large meta-analyses that found climate may have a more variable role in regulating sesquiterpene production (Bourtsoukidis et al. 2024, Bourtsoukidis et al. 2025). Although our sampling methods also may have influenced our characterization of the expression of compounds more susceptible to volatilization like open chain or acyclic monoterpenes, studies have found that the majority of compounds are captured by our extraction process (Clancy et al. 2016). Our results provide initial candidate compounds (and a broader compound class) that might be largely affected by climate change, though additional work is needed to further partition the individual effects of compound changes on phenotypes and fitness in abiotic and biotic environments.

Changes in production of bicyclic monoterpenes likely have consequences for mitigating environmental stressors and there may be an associated adaptive significance to these changes. Bicyclic monoterpenes have been widely studied in plant-herbivore defense and abiotic stress tolerance (Clancy et al. 2016, Zámboriné Németh and Nguyen 2020, Lanier et al. 2023, Rahimova et al. 2024, Shiojiri et al. 2025). Across common tansy metabolomic studies, thujones, sabinenes, and pinenes are generally dominant components (Holopainen et al. 1987, Keskitalo et al. 2001, Zámboriné Németh and Nguyen 2020). Increased production (or proportion of specific compounds within chemotypes) has been previously shown to decrease herbivory directly via increased deterrence (insects: beta-thujone: Kleine and Müller 2011, Ojeda Prieto et al. 2025; alpha-pinene: Erbilgin and Raffa 2000, mammals: alpha/beta-thujone, alpha/beta-pinene and sabinene: Vourc’h et al. 2002) and indirectly via increased attraction of insect predators (alpha-thujone: Senft et al. 2019; alpha-pinene: Mumm and Hilker 2006, Dudareva et al. 2006). Similarly, abiotic stress has also been previously shown to influence bicyclic monoterpene production. While drought has been shown to elevate monoterpene production (e.g. Jamieson et al. 2017; Ramezani et al 2020, Soltanbeigi et al. 2021), it can also nonlinearly affect the production of compounds in the thujone, pinene, and sabinene families (e.g. production increased in moderate but decreased in severe drought; Tan and Gören 2024). The rigid conformation of bicyclic monoterpenes confers high structural specificity for target-receptor binding (Höld et al. 2000, Zielińska-Błajet and Feder-Kubis 2020) and enables membrane-associated antioxidant activities (Soares-Castro et al. 2021). This may help ameliorate membrane disfunction and oxidative damage during environmental stress (Vickers et al. 2008). The changes that we found were likely modulated by a complex interaction between biotic defense, especially in areas where herbivory is greatest, and differing levels of abiotic environmental stress.

### Implications for invasive range dynamics under climate change

Our finding that responses to climate change depend upon source population has important implications for the development of models that forecast geographic range shifts with climate change. Species distribution models (SDMs) typically assume that there is no population differentiation across species’ ranges and that environmental responses are invariant across populations. One reason for this assumption is that data from common garden experiments are rarely available and can be complex to incorporate into distribution models. Nevertheless, evidence is accumulating that predictions of geographic range shifts may be substantially different when evolution is taken into account. Our ongoing work is examining the consequences of incorporating vs. ignoring population differentiation in ecologically-important traits (e.g. terpene profiles and chemotypes) and fitness based on our transplant experiments.

Due to the assumption that geographic ranges are largely controlled by abiotic factors, range shift predictions (using SDMs) also often fail to incorporate biotic interactions (Wisz et al. 2013). At the same time, there is increasing evidence that biotic interactions can be pivotal in modulating stable range limits (Benning et al. 2019) and might influence the pace and/or extent of range shifts with climate change (HilleRisLambers et al. 2013). In this study, the variation in herbivore pressure (and resistance to herbivory) across the invaded range suggests that these plant-herbivore interactions may have the potential influence range dynamics of invasive species. Further, climate change treatments also caused an increased in herbivory (for some chemotypes) in this study, which suggests that variation in herbivory may continue to shift (for certain chemotypes) as the climate changes. As a result, omitting biotic interactions from range shift predictions may weaken forecasts of geographic range shifts in the era of climate change.

## REFERENCES

Adams, J. M., & Zhang, Y. (2009). Is there more insect folivory in warmer temperate climates? A latitudinal comparison of insect folivory in eastern North America. Journal of Ecology, 97(5), 933–940.

Afroz, M., Rahman, M. M., & Amin, M. R. (2021). Insect Plant Interaction with Reference to Secondary Metabolites: A Review. Agricultural Reviews, 42(4), 427.

Akbar, R., Sun, J., Bo, Y., Khattak, W. A., Khan, A. A., Jin, C., … & Du, D. (2024). Understanding the influence of secondary metabolites in plant invasion strategies: A comprehensive review. Plants, 13(22), 3162.

Anstett, D. N., Nunes, K. A., Baskett, C., & Kotanen, P. M. (2016). Sources of controversy surrounding latitudinal patterns in herbivory and defense. Trends in Ecology & Evolution, 31(10), 789–802.

Arias, P. A., Bellouin, N., Coppola, E., Jones, R. G., Krinner, G., Marotzke, J., … & Zhang, X. (2021). Technical summary.

Aslani, Z., Hassani, A., Mandoulakani, B. A., Barin, M., & Maleki, R. (2023). Effect of drought stress and inoculation treatments on nutrient uptake, essential oil and expression of genes related to monoterpenes in sage (Salvia officinalis). Scientia Horticulturae, 309, 111610.

Benedek, K., Bálint, J., Salamon, R. V., Kovács, E., Ábrahám, B., Fazakas, C., … & Balog, A. (2015). Chemotype of tansy (Tanacetum vulgare L.) determines aphid genotype and its associated predator system. Biological Journal of the Linnean Society, 114(3), 709–719.

Benning, J. W., Eckhart, V. M., Geber, M. A., & Moeller, D. A. (2019). Biotic interactions contribute to the geographic range limit of an annual plant: herbivory and phenology mediate fitness beyond a range margin. The American Naturalist, 193(6), 786–797.

Berini, J. L., Brockman, S. A., Hegeman, A. D., Reich, P. B., Muthukrishnan, R., Montgomery, R. A., & Forester, J. D. (2018). Combinations of abiotic factors differentially alter production of plant secondary metabolites in five woody plant species in the boreal-temperate transition zone. Frontiers in plant science, 9, 1257.

Boncan, D. A. T., Tsang, S. S., Li, C., Lee, I. H., Lam, H. M., Chan, T. F., & Hui, J. H. (2020). Terpenes and terpenoids in plants: Interactions with environment and insects. International Journal of Molecular Sciences, 21(19), 7382.

Bourtsoukidis, E., Guenther, A., Wang, H., Economou, T., Lazoglou, G., Christodoulou, A., … & Peñuelas, J. (2025). Environmental change is reshaping the temperature sensitivity of sesquiterpene emissions and their atmospheric impacts. Global change biology, 31(6), e70258.

Bourtsoukidis, E., Pozzer, A., Williams, J., Makowski, D., Peñuelas, J., Matthaios, V. N., … & Sciare, J. (2024). High temperature sensitivity of monoterpene emissions from global vegetation. Communications Earth & Environment, 5(1), 23.

Briscoe Runquist, R., Benning, J. W., & Moeller, D. A. (2026). Expansion load reduces fitness at the range margin of an invasive plant. bioRxiv, 2026-07.

Briscoe Runquist, R., & Moeller, D. A. (2024). Isolation by environment and its consequences for range shifts with global change: Landscape genomics of the invasive plant common tansy. Molecular ecology, 33(16), e17462.

Byrd, D. W., McArthur, E. D., Wang, H., Graham, J. H., & Freeman, D. C. (1999). Narrow hybrid zone between two subspecies of big sagebrush, Artemisia tridentata (Asteraceae). VIII. Spatial and temporal pattern of terpenes. Biochemical Systematics and Ecology, 27(1), 11–25.

Cao, P., Liao, Z. Y., Wang, S., Parepa, M., Zhang, L., Bi, J., … & Li, B. (2025). Cross continental variation of herbivore resistance in a global plant invader. Ecography, 2025(7), e07569.

Carley, L. N., Mojica, J. P., Wang, B., Chen, C. Y., Lin, Y. P., Prasad, K. V., … & Mitchell-Olds, T. (2021). Ecological factors influence balancing selection on leaf chemical profiles of a wildflower. Nature ecology & evolution, 5(8), 1135–1144.

Castells, E., & Sanchez Martinez, P. (2025). The influence of aridity on plant intraspecific chemical diversity supports adaptive differentiation and convergent evolution. Plant Biology, 27(5), 761–772.

Chatterjee, A., & Pal, H. (2023). Terpenoids in Plant Tolerance Against Different Environmental Stress. In *Biology and* Biotechnology of Environmental Stress Tolerance in Plants (pp. 137–161). Apple Academic Press.

Charrad, M., Ghazzali, N., Boiteau, V., & Niknafs, A. (2014). NbClust: an R package for determining the relevant number of clusters in a data set. Journal of statistical software, 61, 1–36.

Chen, L., & Ford, T. W. (2023). Future changes in the transitions of monthly to seasonal precipitation extremes over the Midwest in Coupled Model Intercomparison Project Phase 6 models. International Journal of Climatology, 43(1), 255–274.

Clancy, M. V., Zytynska, S. E., Senft, M., Weisser, W. W., & Schnitzler, J. P. (2016). Chemotypic variation in terpenes emitted from storage pools influences early aphid colonisation on tansy. Scientific reports, 6(1), 38087.

Clasen, B., Moss, N., Chandler, M., & Smith, A. (2011). A preliminary genetic structure study of the non-native weed, common tansy (Tanacetum vulgare). Canadian Journal of Plant Science, 91(4), 717–723.

Colautti, R. I., Alexander, J. M., Dlugosch, K. M., Keller, S. R., & Sultan, S. E. (2017). Invasions and extinctions through the looking glass of evolutionary ecology. Philosophical Transactions of the Royal Society B: Biological Sciences, 372(1712), 20160031.

Comstock, R. E., & Robinson, H. F. (1948). The components of genetic variance in populations of biparental progenies and their use in estimating the average degree of dominance. Biometrics, 4(4), 254–266.

Copolovici, L. O., & Niinemets, Ü. (2005). Temperature dependencies of Henry’s law constants and octanol/water partition coefficients for key plant volatile monoterpenoids. Chemosphere, 61(10), 1390–1400.

Degenhardt, J., Köllner, T. G., & Gershenzon, J. (2009). Monoterpene and sesquiterpene synthases and the origin of terpene skeletal diversity in plants. Phytochemistry, 70(15-16), 1621–1637.

Dobhal, P., Purohit, V. K., Chandra, S., Rawat, S., Prasad, P., Bhandari, U., … & Nautiyal, M. C. (2024). Climate-induced changes in essential oil production and terpene composition in alpine aromatic plants. Plant Stress, 12, 100445.

Dudareva, N., Negre, F., Nagegowda, D. A., & Orlova, I. (2006). Plant volatiles: recent advances and future perspectives. Critical reviews in plant sciences, 25(5), 417–440.

Erbilgin, N., & Raffa, K. F. (2000). Opposing effects of host monoterpenes on responses by two sympatric species of bark beetles to their aggregation pheromones. Journal of Chemical Ecology, 26(11), 2527–2548.

Felisberto, J. S., Machado, D. B., Assunção, J. A., Massau, S. A., Queiroz, G. A. D., Guimarães, E. F., … & Moreira, D. D. L. (2024). Spatio-temporal variations of volatile metabolites as an eco-physiological response of a native species in the tropical forest. Plants, 13(18), 2599.

Gallagher, R. V., Beaumont, L. J., Hughes, L., & Leishman, M. R. (2010). Evidence for climatic niche and biome shifts between native and novel ranges in plant species introduced to Australia. Journal of ecology, 98(4), 790–799.

Gargallo-Garriga, A., Ayala-Roque, M., Sardans, J., Bartrons, M., Granda, V., Sigurdsson, B. D., … & Peñuelas, J. (2017). Impact of soil warming on the plant metabolome of Icelandic grasslands. Metabolites, 7(3), 44.

Gassmann, A., Schwann, A., Mosyakin, A., Wolf, V., Jović, J., & Toševski, I. (2010). Biological control of common tansy, Tanacetum vulgare (p. 43). Annual Report.

Gershenzon, J. (2017). The cost of plant chemical defense against herbivory: a biochemical perspective. In Insect-plant interactions *(1993)* (pp. 105–176). CRC Press.).

Gitonga, H. W., Kyamanywa, S., Arusei, P., Lukanda, M. M., Edema, R., & Dramadri, I. O. (2022). Genotype × Environment interaction influence secondary metabolite in cowpea infested by flower bud thrips. Agronomy, 12(12), 3210.

Hamann, E., Blevins, C., Franks, S. J., Jameel, M. I., & Anderson, J. T. (2021). Climate change alters plant–herbivore interactions. New Phytologist, 229(4), 1894–1910.

Hargreaves, A. L., Suárez, E., Mehltreter, K., Myers-Smith, I., Vanderplank, S. E., Slinn, H. L., … & Morales M. P. A. (2019). Seed predation increases from the Arctic to the Equator and from high to low elevations. Science Advances, 5(2), eaau4403.

He, Y., Junker, R. R., Xiao, J., Lasky, J. R., Cao, M., Asefa, M., … & Sedio, B. E. (2025). Genetic and environmental drivers of intraspecific variation in foliar metabolites in a tropical tree community. New Phytologist, 246(6), 2551–2564.

He, J., Yao, L., Pecoraro, L., Liu, C., Wang, J., Huang, L., & Gao, W. (2023). Cold stress regulates accumulation of flavonoids and terpenoids in plants by phytohormone, transcription process, functional enzyme, and epigenetics. Critical reviews in biotechnology, 43(5), 680–697.

Herve, M. (2025). _RVAideMemoire: Testing and Plotting Procedures for Biostatistics_. R package version 0.9-83-12, <https://CRAN.R-project.org/package=RVAideMemoire>.

Hijmans, R. (2025). _terra: Spatial Data Analysis_. R package version 1.8-60, <https://CRAN.R-project.org/package=terra>.

Hijmans R (2024). _geosphere: Spherical Trigonometry_. R package version 1.5-20, <https://CRAN.R-project.org/package=geosphere>.

HilleRisLambers, J., Harsch, M. A., Ettinger, A. K., Ford, K. R., & Theobald, E. J. (2013). How will biotic interactions influence climate change–induced range shifts?. Annals of the new York Academy of Sciences, 1297(1), 112–125.

Höld, K. M., Sirisoma, N. S., Ikeda, T., Narahashi, T., & Casida, J. E. (2000). α-Thujone (the active component of absinthe): γ-aminobutyric acid type A receptor modulation and metabolic detoxification. Proceedings of the National Academy of Sciences, 97(8), 3826–3831.

Holopainen, J. K., Himanen, S. J., Yuan, J. S., Chen, F., & Stewart Jr, C. N. (2013). Ecological functions of terpenoids in changing climates. In Natural Products: Phytochemistry, Botany, Metabolism of Alkaloids, Phenolics and Terpenes (pp. 1–44). Berlin, Heidelberg: Springer Berlin Heidelberg.

Holopainen, M., Hiltunen, R., & von Schantz, M. (1987). A study on tansy chemotypes. Planta medica, 53(03), 284–287.

Jamieson, M. A., Burkle, L. A., Manson, J. S., Runyon, J. B., Trowbridge, A. M., & Zientek, J. (2017). Global change effects on plant–insect interactions: the role of phytochemistry. Current opinion in insect science, 23, 70–80.

Jamloki, A., Bhattacharyya, M., Nautiyal, M. C., & Patni, B. (2021). Elucidating the relevance of high temperature and elevated CO2 in plant secondary metabolites (PSMs) production. Heliyon, 7(8).

Jogawat, A., Yadav, B., Chhaya, Lakra, N., Singh, A. K., & Narayan, O. P. (2021). Crosstalk between phytohormones and secondary metabolites in the drought stress tolerance of crop plants: a review. Physiologia Plantarum, 172(2), 1106–1132.

Kaplan, F., Kopka, J., Haskell, D. W., Zhao, W., Schiller, K. C., Gatzke, N., … & Guy, C. L. (2004). Exploring the temperature-stress metabolome of Arabidopsis. Plant physiology, 136(4), 4159–4168.

Kariñho-Betancourt, E. (2018). Plant-herbivore interactions and secondary metabolites of plants: Ecological and evolutionary perspectives. Botanical Sciences, 96(1), 35–51.

Kassambara, A. & Mundt, F. (2020). _factoextra: Extract and Visualize the Results of Multivariate Data Analyses_. R package version 1.0.7, <https://CRAN.R-project.org/package=factoextra>.

Keefover Ring, K. (2022). The chemical biogeography of a widespread aromatic plant species shows both spatial and temporal variation. Ecology and Evolution, 12(9).

Kent, D. R., Lynn, J. S., Pennings, S. C., Souza, L. A., Smith, M. D., & Rudgers, J. A. (2020). Weak latitudinal gradients in insect herbivory for dominant rangeland grasses of North America. Ecology and Evolution, 10(13), 6385–6394.

Kesik, T., Wojciechowska, A. E., & Pitura, K. (2008). Differentiation of morphological traits of common tansy (Tanacetum vulgare l.) originating from different stands. Acta Agrobotanica, 61(2).

Keskitalo, M., Pehu, E., & Simon, J. E. (2001). Variation in volatile compounds from tansy (Tanacetum vulgare L.) related to genetic and morphological differences of genotypes. Biochemical systematics and ecology, 29(3), 267–285.

Kessler, A., & Kalske, A. (2018). Plant secondary metabolite diversity and species interactions. Annual review of ecology, evolution, and systematics, 49(1), 115–138.

Kivimäenpää, M., Riikonen, J., Valolahti, H., Elina, H., Holopainen, J. K., & Holopainen, T. (2022). Effects of elevated ozone and warming on terpenoid emissions and concentrations of Norway spruce depend on needle phenology and age. Tree Physiology, 42(8), 1570–1586.

Kleine, S., & Müller, C. (2011). Intraspecific plant chemical diversity and its relation to herbivory. Oecologia, 166(1), 175–186.

Kohl, S. M., Klein, M. S., Hochrein, J., Oefner, P. J., Spang, R., & Gronwald, W. (2012). State-of-the art data normalization methods improve NMR-based metabolomic analysis. Metabolomics, 8(Suppl 1), 146–160.

Kreuzwieser, J., Meischner, M., Grün, M., Yáñez Serrano, A. M., Fasbender, L., & Werner, C. (2021). Drought affects carbon partitioning into volatile organic compound biosynthesis in Scots pine needles. New Phytologist, 232(5), 1930–1943.

Lanier, E. R., Andersen, T. B., & Hamberger, B. (2023). Plant terpene specialized metabolism: complex networks or simple linear pathways?. The Plant Journal, 114(5), 1178–1201.

Leisso, R., Rudell, D., & Mazzola, M. (2018). Targeted metabolic profiling indicates apple rootstock genotype-specific differences in primary and secondary metabolite production and validate quantitative contribution from vegetative growth. Frontiers in Plant Science, 9, 1336.

Levin, D. A. (1976). The chemical defenses of plants to pathogens and herbivores. Annual review of Ecology and Systematics, 121–159.

Liu, M., Jiang, P., Chase, J. M., & Liu, X. (2024). Global insect herbivory and its response to climate change. Current Biology, 34(12), 2558–2569.

Liu, M., Pan, Y., Pan, X., Sosa, A., Blumenthal, D. M., Van Kleunen, M., & Li, B. (2021). Plant invasion alters latitudinal pattern of plant defense syndromes. Ecology, 102(12), e03511.

Liu, B., Zhang, L., Rusalepp, L., Kaurilind, E., Sulaiman, H. Y., Püssa, T., & Niinemets, Ü. (2021). Heat priming improved heat tolerance of photosynthesis, enhanced terpenoid and benzenoid emission and phenolics accumulation in Achillea millefolium. Plant, Cell & Environment, 44(7), 2365–2385.

Llusià, J., & Peñuelas, J. (1998). Changes in terpene content and emission in potted Mediterranean woody plants under severe drought. Canadian Journal of Botany, 76(8), 1366–1373.

Loreto, F., Förster, A., Dürr, M., Csiky, O., & Seufert, G. (1998). On the monoterpene emission under heat stress and on the increased thermotolerance of leaves of Quercus ilex L. fumigated with selected monoterpenes. *Plant*, Cell & Environment, 21(1), 101–107.

Magnani, R. F., Volpe, H. X. L., Luvizotto, R. A. G., Mulinari, T. A., Agostini, T. T., Bastos, J. K., … & Leal, W. S. (2025). α-Copaene is a potent repellent against the Asian Citrus Psyllid Diaphorina citri. Scientific Reports, 15(1), 3564.

Matesanz, S., Horgan-Kobelski, T., & Sultan, S. E. (2012). Phenotypic plasticity and population differentiation in an ongoing species invasion. PLoS ONE, 7(9): e44955. doi:10.1371/journal.pone.0044955

Mihaliak, C. A., & Lincoln, D. E. (1989). Changes in leaf mono-and sesquiterpene metabolism with nitrate availability and leaf age in Heterotheca subaxillaris. Journal of chemical ecology, 15(5), 1579–1588.

Moles, A. T., Bonser, S. P., Poore, A. G., Wallis, I. R., & Foley, W. J. (2011). Assessing the evidence for latitudinal gradients in plant defence and herbivory. Functional Ecology, 25(2), 380–388.

Mooney, H. A., & Cleland, E. E. (2001). The evolutionary impact of invasive species. Proceedings of the National Academy of Sciences, 98(10), 5446–5451.

Moore, B. D., Andrew, R. L., Külheim, C., & Foley, W. J. (2014). Explaining intraspecific diversity in plant secondary metabolites in an ecological context. New Phytologist, 201(3), 733–750.

Moran, E. V., & Alexander, J. M. (2014). Evolutionary responses to global change: lessons from invasive species. Ecology Letters, 17(5), 637–649.

Müller, C., Dussarrat, T., & van Dam, N. M. (2026). What is a plant chemotype, anyway?. Trends in Plant Science.

Mumm, R., Posthumus, M. A., & Dicke, M. (2008). Significance of terpenoids in induced indirect plant defence against herbivorous arthropods. Plant, Cell & Environment, 31(4), 575–585.

Mumm, R., & Hilker, M. (2006). Direct and indirect chemical defence of pine against folivorous insects. Trends in plant science, 11(7), 351–358.

Nakabayashi, R., Yonekura Sakakibara, K., Urano, K., Suzuki, M., Yamada, Y., Nishizawa, T., … & Saito, K. (2014). Enhancement of oxidative and drought tolerance in Arabidopsis by overaccumulation of antioxidant flavonoids. The Plant Journal, 77(3), 367–379.

Nielsen, J. A., Whigham, P. A., Frew, R. D., Callaway, R. M., & Dickinson, K. J. M. (2014). Invasion essentials: does secondary chemistry plasticity contribute to the invasiveness of Thymus vulgaris L.?. Chemoecology, 24(1), 15–27.

Neilson, E. H., Goodger, J. Q., Woodrow, I. E., & Møller, B. L. (2013). Plant chemical defense: at what cost?. Trends in plant science, 18(5), 250–258.

Newrzella, H., Heinen, R., Villasante, C. M., Zimmer, I., Weber, B., Kary, P., … & Schnitzler, J. P. (2026). Organ specific terpenoid responses in Tanacetum vulgare are chemotype dependent. Plant Biology, 28(5), 1437–1453.

Neuhaus-Harr, A., Ojeda-Prieto, L., Zhang, X., Schnitzler, J. P., Weisser, W. W., & Heinen, R. (2025). Aphid colony size in tansy is affected by plant chemical composition but not by belowground herbivory. Journal of Chemical Ecology, 51(3), 63.

Neuhaus Harr, A., Ojeda Prieto, L., Eilers, E., Müller, C., Weisser, W. W., & Heinen, R. (2024). Chemodiversity affects preference for Tanacetum vulgare chemotypes in two aphid species. Oikos, 2024(3), e10437.

Nomoto, H., Fernández Conradi, P., Kjelsberg, N., Defossez, E., Münzbergová, Z., Glauser, G., & Rasmann, S. (2025). Elevation drives intraspecific metabolomic differentiation in natural and experimental populations. Plant Biology, 27(5), 773–784.

Obata, T., & Fernie, A. R. (2012). The use of metabolomics to dissect plant responses to abiotic stresses. Cellular and Molecular Life Sciences, 69(19), 3225–3243.

Oduor, A. M. (2022). Invasive plant species that experience lower herbivory pressure may evolve lower diversities of chemical defense compounds in the exotic range. American Journal of Botany, 109(9), 1382–1393.

Ojeda Prieto, L., Moreno, E. L., Heinen, R., & Weisser, W. W. (2025). Intraspecific plant chemodiversity at plot level has contrasting effects on arthropod functional groups. Functional Ecology, 39(12), 3732–3750.

Ojha, M., Naidu, D. G., & Bagchi, S. (2022). Meta analysis of induced anti herbivore defence traits in plants from 647 manipulative experiments with natural and simulated herbivory. Journal of Ecology, 110(4), 799–816.

Okamura, Y., Sawada, Y., Hirai, M. Y., & Murakami, M. (2016). Effects of different secondary metabolite profiles in plant defense syndromes on specialist and generalist herbivores. Entomological Science, 19(2), 97–103.

Oksanen J, Simpson G, Blanchet F, Kindt R, Legendre P, Minchin P, O’Hara R, Solymos P, Stevens M, Szoecs E, Wagner H, Barbour M, Bedward M, Bolker B, Borcard D, Borman T, Carvalho G, Chirico M, De Caceres M, Durand S, Evangelista H, FitzJohn R, Friendly M, Furneaux B, Hannigan G, Hill M, Lahti L, Martino C, McGlinn D, Ouellette M, Ribeiro Cunha E, Smith T, Stier A, Ter Braak C, Weedon J (2025). _vegan: Community Ecology Package_. R package version 2.7-1, <https://CRAN.R-project.org/package=vegan>.

Ormeño, E., Baldy, V., Ballini, C., & Fernandez, C. (2008). Production and diversity of volatile terpenes from plants on calcareous and siliceous soils: effect of soil nutrients. Journal of chemical ecology, 34(9), 1219–1229.

Parida, A. K., Panda, A., & Rangani, J. (2018). Metabolomics-guided elucidation of abiotic stress tolerance mechanisms in plants. In Plant metabolites and regulation under environmental stress (pp. 89-131). Academic Press.

Peñuelas, J., Sardans, J., Llusia, J., Owen, S. M., Silva, J., & Niinemets, Ü. (2010). Higher allocation to low cost chemical defenses in invasive species of Hawaii. Journal of Chemical Ecology, 36(11), 1255–1270.

Petrén, H., Köllner, T.G., and Junker, R.R. (2023). Quantifying chemodiversity considering biochemical and structural properties of compounds with the R package chemodiv. New Phytologist 237: 2478–2492.

Pichersky, E., & Raguso, R. A. (2018). Why do plants produce so many terpenoid compounds?. New Phytologist, 220(3), 692–702.

Pratt, J. D., Keefover Ring, K., Liu, L. Y., & Mooney, K. A. (2014). Genetically based latitudinal variation in Artemisia californica secondary chemistry. Oikos, 123(8), 953–963.

Prentis, P. J., Wilson, J. R., Dormontt, E. E., Richardson, D. M., & Lowe, A. J. (2008). Adaptive evolution in invasive species. Trends in plant science, 13(6), 288–294.

Rahimova, H., Neuhaus Harr, A., Clancy, M. V., Guo, Y., Junker, R. R., Ojeda Prieto, L., … & Schnitzler, J. P. (2024). Geographic distribution of terpenoid chemotypes in Tanacetum vulgare mediates tansy aphid occurrence but not abundance. Oikos, 2024(7), e10320.

Ramezani, S., Abbasi, A., Sobhanverdi, S., Shojaeiyan, A., & Ahmadi, N. (2020). The effects of water deficit on the expression of monoterpene synthases and essential oils composition in Salvia ecotypes. Physiology and Molecular Biology of Plants, 26(11), 2199–2207.

Rasmann, S., & Agrawal, A. A. (2011). Latitudinal patterns in plant defense: evolution of cardenolides, their toxicity and induction following herbivory. Ecology Letters, 14(5), 476–483.

Redman, A. M., Cipollini Jr, D. F., & Schultz, J. C. (2001). Fitness costs of jasmonic acid-induced defense in tomato, Lycopersicon esculentum. Oecologia, 126(3), 380–385.

Rohloff, J., Mordal, R., & Dragland, S. (2004). Chemotypical variation of tansy (Tanacetum vulgare L.) from 40 different locations in Norway. Journal of Agricultural and Food Chemistry, 52(6), 1742–1748.

Romero, H., Pott, D. M., Vallarino, J. G., & Osorio, S. (2021). Metabolomics-based evaluation of crop quality changes as a consequence of climate change. Metabolites, 11(7), 461.

Rowen, E., & Kaplan, I. (2016). Eco evolutionary factors drive induced plant volatiles: a meta analysis. New Phytologist, 210(1), 284–294.

Rudgers, J. A., Luketich, A., Bacigalupa, M., Baur, L. E., Collins, S. L., Hall, K. M., … & Zhou, Y. (2023). Infrastructure to factorially manipulate the mean and variance of precipitation in the field. Ecosphere, 14(7), e4603.

Saddique, M., Kamran, M., & Shahbaz, M. (2018). Differential responses of plants to biotic stress and the role of metabolites. In Plant metabolites and regulation under environmental stress (pp. 69–87). Academic Press.

Sampaio, B. L., Edrada-Ebel, R., & Da Costa, F. B. (2016). Effect of the environment on the secondary metabolic profile of Tithonia diversifolia: a model for environmental metabolomics of plants. Scientific reports, 6(1), 29265.

Sanchez, D. H., Siahpoosh, M. R., Roessner, U., Udvardi, M., & Kopka, J. (2008). Plant metabolomics reveals conserved and divergent metabolic responses to salinity. Physiologia plantarum, 132(2), 209–219.

Sardans, J., Gargallo-Garriga, A., Urban, O., Klem, K., Walker, T. W., Holub, P., … & Peñuelas, J. (2020). Ecometabolomics for a better understanding of plant responses and acclimation to abiotic factors linked to global change. Metabolites, 10(6), 239.

Sardans, J., Penuelas, J., & Rivas-Ubach, A. (2011). Ecological metabolomics: overview of current developments and future challenges. Chemoecology, 21(4), 191–225.

Schemske, D. W., Mittelbach, G. G., Cornell, H. V., Sobel, J. M., & Roy, K. (2009). Is there a latitudinal gradient in the importance of biotic interactions?. Annu. Rev. Ecol. Evol. Syst., 40(1), 245–269.

Senft, M., Clancy, M. V., Weisser, W. W., Schnitzler, J. P., & Zytynska, S. E. (2019). Additive effects of plant chemotype, mutualistic ants and predators on aphid performance and survival. Functional Ecology, 33(1), 139–151.

Shiojiri, K., Kagiya, S., Hagiwara, T., & Karban, R. (2025). Long-term neighbor identity influences effectiveness of sagebrush responses to damage-induced volatiles of those neighbors. Scientific Reports, 15(1), 40056.

Soares-Castro, P., Soares, F., & Santos, P. M. (2020). Current advances in the bacterial toolbox for the biotechnological production of monoterpene-based aroma compounds. Molecules, 26(1), 91.

Soltanbeigi, A., Yıldız, M., Dıraman, H., Terzi, H., Sakartepe, E., & Yıldız, E. (2021). Growth responses and essential oil profile of Salvia officinalis L. influenced by water deficit and various nutrient sources in the greenhouse. Saudi Journal of Biological Sciences, 28(12), 7327–7335.

Strauss, S. Y., & Zangerl, A. R. (2002). Plant-insect interactions in terrestrial ecosystems. Plant-animal interactions: an evolutionary approach, 2002, 77–106.

Sun, Y., & Fernie, A. R. (2024). Plant secondary metabolism in a fluctuating world: climate change perspectives. Trends in Plant Science, 29(5), 560–571.

Sun, J., & Xia, Y. (2024). Pretreating and normalizing metabolomics data for statistical analysis. Genes & Diseases, 11(3), 100979.

Sun, Y., Züst, T., Silvestro, D., Erb, M., Bossdorf, O., Mateo, P., … & Müller Schärer, H. (2022). Climate warming can reduce biocontrol efficacy and promote plant invasion due to both genetic and transient metabolomic changes. Ecology Letters, 25(6), 1387–1400.

Tan, U., & Gören, H. K. (2024). Comprehensive evaluation of drought stress on medicinal plants: a meta-analysis. PeerJ, 12, e17801.

Unsicker, Sybille B., Grit Kunert, and Jonathan Gershenzon. (2009). Protective perfumes: the role of vegetative volatiles in plant defense against herbivores. Current opinion in plant biology 12.4, 479–485.

van Boheemen, L. A., Atwater, D. Z., & Hodgins, K. A. (2019). Rapid and repeated local adaptation to climate in an invasive plant. New Phytologist, 222(1), 614–627.

Večeřová, K., Klem, K., Veselá, B., Holub, P., Grace, J., & Urban, O. (2021). Combined effect of altitude, season and light on the accumulation of extractable terpenes in Norway spruce needles. Forests, 12(12), 1737.

Vickers, C. E., Gershenzon, J., Lerdau, M. T., & Loreto, F. (2009). A unified mechanism of action for volatile isoprenoids in plant abiotic stress. Nature chemical biology, 5(5), 283–291.

Vourc’h, G., De Garine-Wichatitsky, M., Labbé, A., Rosolowski, D., Martin, J. L., & Fritz, H. (2002). Monoterpene effect on feeding choice by deer. Journal of chemical ecology, 28(12), 2411–2427.

Wada, H., & Sewall, K. B. (2014). Introduction to the symposium—uniting evolutionary and physiological approaches to understanding phenotypic plasticity. American Zoologist, 54(5), 774–782.

Whitehead, S. R., Bass, E., Corrigan, A., Kessler, A., & Poveda, K. (2021). Interaction diversity explains the maintenance of phytochemical diversity. Ecology Letters, 24(6), 1205–1214.

Wittmann, M. J., & Bräutigam, A. (2025). How does plant chemodiversity evolve? Testing five hypotheses in one population genetic model. New Phytologist, 245(3), 1302–1314.

Wisz, M. S., Pottier, J., Kissling, W. D., Pellissier, L., Lenoir, J., Damgaard, C. F., … & Svenning, J. C. (2013). The role of biotic interactions in shaping distributions and realised assemblages of species: implications for species distribution modelling. Biological reviews, 88(1), 15–30.

Wolf, V. C., Berger, U., Gassmann, A., & Müller, C. (2011). High chemical diversity of a plant species is accompanied by increased chemical defence in invasive populations. Biological Invasions, 13(9), 2091–2102.

Xiao, L., Ding, J., Zhang, J., Huang, W., & Siemann, E. (2020). Chemical responses of an invasive plant to herbivory and abiotic environments reveal a novel invasion mechanism. Science of the Total Environment, 741, 140452.

Yang, L., Wen, K. S., Ruan, X., Zhao, Y. X., Wei, F., & Wang, Q. (2018). Response of plant secondary metabolites to environmental factors. Molecules, 23(4), 762.

Yuan, J. S., Himanen, S. J., Holopainen, J. K., Chen, F., & Stewart, C. N. (2009). Smelling global climate change: mitigation of function for plant volatile organic compounds. Trends in Ecology & Evolution, 24(6), 323–331.

Zámboriné Németh, É., & Thi Nguyen, H. (2020). Thujone, a widely debated volatile compound: What do we know about it?. Phytochemistry reviews, 19(2), 405–423.

Zhao, M., Zhang, N., Gao, T., Jin, J., Jing, T., Wang, J., … & Song, C. (2020). Sesquiterpene glucosylation mediated by glucosyltransferase UGT91Q2 is involved in the modulation of cold stress tolerance in tea plants. New phytologist, 226(2), 362–372.

Zhou, L., Yin, W., & Ding, J. (2024). Trade-offs between chemical resistance to herbivory and responses to abiotic stresses in invasive plants. Journal of Plant Ecology, 17(2), rtae007.

Ziaja, D., & Müller, C. (2025). Intraspecific and intra individual chemodiversity and phenotypic integration of terpenes across plant parts and development stages in an aromatic plant. Plant Biology, 27(5), 637–650.

Zielińska-Błajet, M., & Feder-Kubis, J. (2020). Monoterpenes and their derivatives—Recent development in biological and medical applications. International Journal of Molecular Sciences, 21(19), 7078.

Züst, T., & Agrawal, A. A. (2017). Trade-offs between plant growth and defense against insect herbivory: an emerging mechanistic synthesis. Annual review of plant biology, 68(1), 513–534.

