## Supplementary Information for "Climate change effects on terpene expression differ among populations and have consequences for herbivory in an invasive plant"

**Supplemental Information**

**Site, population, and treatment effects on terpene composition (Jaccard dissimilarity)**

***Methods.*** To evaluate spatial (2025 terpenes) and temporal (combined 2024 and 2025 terpenes) compositional differences in terpene production, we used a PERMANOVA with Jaccard dissimilarity (on unmodified peak-normalized area converted to binary data) and 999 permutations using the vegan package (Oksanen et al. 2025). As in the main text, we evaluated significance using type II tests with the RVAideMemoire wrapper package (Herve 2025). We included the fixed effects of source population, transplant site, treatment (control-control or elevated temperature-reduced precipitation), block nested within transplant site, and all two- and three-way interactions. We used the *pairwise.adonis* function in the pairwiseAdonis package (Martinez 2017) to perform post hoc pairwise PERMANOVAs with a correction for multiple comparisons using the Benjamini-Hochberg method. Following PERMANOVAs, we evaluated dispersion of individuals around the group centroids as a validation metric using vegan (*betadisper* function).

To better characterize differences in terpene composition among groups, we examined the relative frequency of compounds. To do so, we used *decostand* function in vegan to convert the binary data into relative frequencies of compounds (proportion of individuals in a particular group) and followed by *simper* function to quantify differences within and between sites (999 permutations). We also used a custom Benjamini-Hochberg correction to identify compounds that strongly contributed to overall compositional dissimilarity between groups.

***Results. Site, population, and treatment effects on terpene composition* –** Variation among transplant site environments influenced terpene composition (Jaccard) (PseudoF_3,311_ 3.46, p < 0.001, Table SA). More specifically, terpene composition of plants in the NE and SW sites both differed from the NW and SE sites but not from each other (pairwise comparisons with NW and SE: all adjusted p-value < 0.05, Table SB). We found no effect of source population or climate change treatment on terpene composition or evidence of any interactions, suggesting that transplant site primarily determined terpene composition. Dispersion of terpene composition among individuals was lower at SE than the other sites (PseudoF_3,308_ = 7.90, p < 0.001).

Our comparison of terpene composition (Jaccard) among sites (mean of individual frequencies) showed that compositional differences were often complex and did not simply reflect site differences in climate (temperature or precipitation). For example, the NE and SW sites had a lower average frequency of isophytol than the NW and SE sites (42-47% cumulative contribution, p-adjusted = 0.014, Table SC). Similarly, the NE and SW sites had a lower frequency of three compounds compared to the SE site (7-methoxycoumarin, car-3-en-5-one, germacrene D-4-ol; all p-adjusted < 0.05, Table S5; all p-adjusted < 0.05, Table SC) and one that was in substantially higher frequency than the NE site (phytol). Finally, the two southern sites differed from one another by one compound (decreased caryophyllene oxide; p-adjusted = 0.014, Table S5).

***Temporal variation in terpene composition –*** In an analysis that combined both years of data for the NW and NE sites, terpene composition differed between years (PseudoF_1,144_ = 15.4, p < 0.001, Table S9), sites (PseudoF_1,144_ = 6.17, p < 0.001), and populations (PseudoF_9,144_ = 15.4, p < 0.001). Importantly, the magnitude of differences between years depended upon population and site (population × site: PseudoF_9,144_ = 1.61, p = 0.081; population × year: PseudoF_9,144_ = 1.74, p < 0.05, Table SD).

For individual compounds, our results indicate that certain terpenes had increased relative frequency of expression in 2024 than 2025. At the NW site, eight compounds that were expressed in 2024 were absent in 2025. At the NE site, four compounds were only expressed in 2024 (sabinene, gamma-muurolene, two neophytadiene isomers) versus one compound only expressed in 2025 (eugenol). Multiple comparisons further demonstrated that relative frequency of compounds differed between most populations × year combinations (p-adjusted < 0.05).

**Additional patterns in terpene expression (non-significant after p-value adjustments)**

***Methods.*** Refer to main text.

***Results.*** We also found tentative evidence consistent with differential terpene expression among source populations. For example, one southern population (WIN020) tended to have a relatively decreased abundance of chrysanthenone compared to two other southern populations (MUR002 and WAB006) (lfc = -4.35 to -4.49, both p < 0.01, both p-adjusted < 0.090). Five additional compounds tended to be less abundant in the southernmost populations (MUR002, WIN020, LINC004, WAB006) compared to some central and northern populations (alpha-thujone, beta-pinene, coumarin, isogermacrene-D, pinocarvone); however, they were not statistically significant after p-value adjustments (total of 14 population-treatment pairs, p < 0.05, p-adjusted > 0.1).

Individuals in the SE site tended to have higher neophytadiene expression compared to the NE site (lfc = 1.06, p = 0.011, p-adjusted = 0.099). We also detected six additional compounds that were differentially expressed among sites (i.e., increased relative abundance in the NE site) that were not statistically significant after p-value adjustments (lfc: -1.30 to 1.06, p < 0.05, p-adjusted > 0.1; Fig. S1). Patterns indicated that central populations (i.e., populations from centermost latitudes) had greater expression of one compound (beta-pinene) in the ambient treatment compared to southern populations (i.e., populations from lower latitudes) in either treatment, but this difference was not statistically significant after p-value adjustments (greater abundance in two of three central-southern population-treatment pairs; p < 0.05, p-adjusted > 0.1).

**Table SA. Terpene composition PERMANOVA results.** PERMANOVA of compounds using a Jaccard dissimilarity matrix evaluated with “Type II” sum of squares. Significant P-values are bolded and denoted as follows: +*P* < 0.1, **P* < 0.05, ***P* < 0.01, ****P* < 0.001.

|  | **Sum Sq** | **Mean Sq** | **Df** | **F** | **P-value** |
| --- | --- | --- | --- | --- | --- |
| Transplant site | 1.251 | 0.139 | 3 | 3.459 | **0.001***** |
| Source population | 0.041 | 0.041 | 9 | 1.027 | 0.401 |
| Treatment | 0.045 | 0.045 | 1 | 1.115 | 0.332 |
| Transplant site × source population | 0.038 | 0.013 | 27 | 0.317 | 0.998 |
| Transplant site × treatment | 0.439 | 0.049 | 3 | 1.215 | 0.152 |
| Source population × treatment | 0.132 | 0.033 | 9 | 0.822 | 0.719 |
| Transplant site × block | 0.134 | 0.033 | 4 | 0.832 | 0.694 |
| Transplant site × source population × treatment | 1.061 | 0.039 | 27 | 0.978 | 0.572 |
| Residual | 9.159 | 0.04 | 228 | NA | NA |
| Total | 14.426 | 0.046 | 311 | NA | NA |

**Table SB. Pairwise comparison of terpene compositional differences in transplant sites.** Pairwise PERMANOVA comparison of compounds using a Jaccard dissimilarity matrix evaluated with “Type II” sum of squares.

| **Pairs** | **Sum Sq** | **F** | **R2** | **P-value** | **P-adj** | **sig** |
| --- | --- | --- | --- | --- | --- | --- |
| NE vs SW | 0.0548 | 1.0796 | 0.0070 | 0.357 | 1 |  |
| NE vs NW | 0.3044 | 6.7180 | 0.0423 | 0.001 | 0.006 | * |
| NE vs SE | 0.3312 | 7.8457 | 0.0488 | 0.001 | 0.006 | * |
| SW vs NW | 0.2500 | 5.3202 | 0.0332 | 0.002 | 0.012 | . |
| SW vs SE | 0.3494 | 7.9536 | 0.0485 | 0.001 | 0.006 | * |
| NW vs SE | 0.0796 | 2.0640 | 0.0131 | 0.056 | 0.336 |  |

**Table SC. Differences in average compound frequency between transplant sites.** Summarized output from the *simper* function in vegan. The “Avg” is the raw absolute average contribution to between-group dissimilarity, “Prop” is the weighed cumulative sum expressed as a proportion, “Sites (A_B)” is the focal pairwise comparison between sites, “FreqA” is the average frequency of the first site listed in the comparison column, “FreqB” is the average frequency of the second site listed in the comparison column, “Direction” summarizes FreqA and FreqB by denoting whether the first site has an increased or decreased compound frequency, and “Rank” is the rank contribution of each compound to the dissimilarity matrix. Note that (E,E)-7,11,15-Trimethyl-3-methylene-hexadeca-1,6,10,14-tetraene and Bicyclosesquiphellandrene were shortened in the table to “Trimethyl” and “Bicyclosesq” to increase table readability.

| **Compound** | **Avg** | **Prop** | **Sites (A_B)** | **FreqA** | **FreqB** | **Direction** | **Rank** | **P-value** | **P-adj** |
| --- | --- | --- | --- | --- | --- | --- | --- | --- | --- |
| Trimethyl | 0.007 | 0.484 | NE_SW | 0.75 | 0.633 | increased | 13 | 0.001 | 0.0144 |
| Trimethyl | 0.007 | 0.733 | SW_NW | 0.633 | 0.821 | decreased | 20 | 0.001 | 0.0144 |
| 7-Methoxycoumarin | 0.008 | 0.088 | NE_SE | 0.421 | 0.709 | decreased | 2 | 0.003 | 0.0311 |
| 7-Methoxycoumarin | 0.009 | 0.086 | SW_SE | 0.38 | 0.709 | decreased | 2 | 0.001 | 0.0144 |
| alpha-Copaene | 0 | 0.994 | NE_SW | 0.987 | 1 | decreased | 41 | 0.001 | 0.0144 |
| alpha-Terpineol | 0.008 | 0.079 | NE_SW | 0.579 | 0.481 | increased | 2 | 0.001 | 0.0144 |
| alpha-Thujene | 0.007 | 0.618 | NE_SW | 0.789 | 0.684 | increased | 17 | 0.003 | 0.0311 |
| alpha-Thujone | 0.008 | 0.493 | SW_NW | 0.696 | 0.564 | increased | 13 | 0.002 | 0.0235 |
| beta-Bourbonene | 0.002 | 0.963 | NE_SW | 0.934 | 0.962 | decreased | 34 | 0.002 | 0.0235 |
| beta-Copaene | 0 | 0.995 | NE_SW | 0.987 | 1 | decreased | 42 | 0.001 | 0.0144 |
| Bicyclosesq | 0.004 | 0.893 | NE_SW | 0.882 | 0.886 | decreased | 28 | 0.004 | 0.0384 |
| Camphene | 0.008 | 0.233 | NE_SW | 0.566 | 0.633 | decreased | 6 | 0.001 | 0.0144 |
| Car-3-en-5-one | 0.008 | 0.13 | NE_SE | 0.276 | 0.582 | decreased | 3 | 0.001 | 0.0144 |
| Car-3-en-5-one | 0.008 | 0.272 | SW_NW | 0.253 | 0.5 | decreased | 7 | 0.006 | 0.0457 |
| Car-3-en-5-one | 0.008 | 0.127 | SW_SE | 0.253 | 0.582 | decreased | 3 | 0.001 | 0.0144 |
| Caryophyllene | 0 | 0.997 | NE_SW | 0.987 | 1 | decreased | 43 | 0.001 | 0.0144 |
| Caryophyllene oxide | 0.008 | 0.04 | NE_SW | 0.605 | 0.468 | increased | 1 | 0.001 | 0.0144 |
| Caryophyllene oxide | 0.008 | 0.081 | SW_NW | 0.468 | 0.641 | decreased | 2 | 0.003 | 0.0311 |
| Caryophyllene oxide | 0.008 | 0.167 | SW_SE | 0.468 | 0.747 | decreased | 4 | 0.002 | 0.0235 |
| Chrysanthenone | 0.007 | 0.681 | NE_SW | 0.711 | 0.747 | decreased | 19 | 0.002 | 0.0235 |
| cis-Cubenol | 0.008 | 0.157 | NE_SW | 0.487 | 0.519 | decreased | 4 | 0.005 | 0.0418 |
| cis-Sabinol | 0.008 | 0.309 | SW_NW | 0.608 | 0.564 | increased | 8 | 0.006 | 0.0457 |
| endo-Borneol | 0.008 | 0.118 | NE_SW | 0.487 | 0.557 | decreased | 3 | 0.004 | 0.0384 |
| Eugenol | 0 | 1 | NE_SW | 0.987 | 1 | decreased | 46 | 0.001 | 0.0144 |
| gamma-Elemene | 0.002 | 0.955 | NE_SW | 0.934 | 0.949 | decreased | 33 | 0.006 | 0.0457 |
| Germacrene D-4-ol | 0.008 | 0.248 | NE_SE | 0.25 | 0.519 | decreased | 6 | 0.005 | 0.0418 |
| Germacrene D-4-ol | 0.008 | 0.245 | SW_SE | 0.342 | 0.519 | decreased | 6 | 0.005 | 0.0418 |
| Isophytol | 0.009 | 0.043 | NE_NW | 0.382 | 0.769 | decreased | 1 | 0.001 | 0.0144 |
| Isophytol | 0.009 | 0.047 | NE_SE | 0.382 | 0.886 | decreased | 1 | 0.001 | 0.0144 |
| Isophytol | 0.009 | 0.042 | SW_NW | 0.418 | 0.769 | decreased | 1 | 0.001 | 0.0144 |
| Isophytol | 0.009 | 0.044 | SW_SE | 0.418 | 0.886 | decreased | 1 | 0.001 | 0.0144 |
| Neophytadiene (Isomer #2) | 0 | 0.998 | NE_SW | 0.987 | 1 | decreased | 44 | 0.001 | 0.0144 |
| Phytol | 0.008 | 0.195 | NE_SW | 0.526 | 0.544 | decreased | 5 | 0.001 | 0.0144 |
| Phytol | 0.008 | 0.273 | NE_NW | 0.526 | 0.731 | decreased | 7 | 0.005 | 0.0418 |

**Table SD. PERMANOVA of terpene composition between years.** PERMANOVA of compounds using a Jaccard dissimilarity matrix evaluated with “Type II” sum of squares. Significant P-values are bolded and denoted as follows: +*P* < 0.1, **P* < 0.05, ***P* < 0.01, ****P* < 0.001.

|  | Sum Sq | Mean Sq | Df | F | P-value |
| --- | --- | --- | --- | --- | --- |
| Transplant site | 11.274 | 1.253 | 1 | 6.173 | **0.001***** |
| Source population | 3.12 | 3.12 | 9 | 15.377 | **0.001***** |
| Year | 3.131 | 3.131 | 1 | 15.43 | **0.001***** |
| Transplant site × source population | 0.332 | 0.332 | 9 | 1.635 | *0.077+* |
| Transplant site × year | 2.947 | 0.327 | 1 | 1.614 | **0.001**** |
| Source population × year | 0.705 | 0.352 | 9 | 1.736 | **0.016*** |
| Transplant site × block | 0.692 | 0.346 | 2 | 1.705 | **0.024*** |
| Residual | 29.219 | 0.203 | 144 | NA | NA |
| Total | 50.639 | 0.288 | 176 | NA | NA |

**Table S1. Locations of source populations.** Regions were determined a priori to experiment onset (BIG002 considered SW population due to region ecological overlap).

| **Population** | **Region** | **Latitude** | **Longitude** |
| --- | --- | --- | --- |
| MARS001 | NW | 48.2658 | -95.9329 |
| PEN002 | NW | 48.02011 | -95.7002 |
| STL035 | NE | 47.90207 | -91.8765 |
| COOK001.5 | NE | 47.77406 | -90.2431 |
| STL037 | NE | 47.68881 | -91.6405 |
| BIG002 | SW | 45.34491 | -96.4009 |
| WAB006 | SE | 44.35179 | -92.3756 |
| LINC004 | SW | 44.2616 | -96.2788 |
| WIN020 | SE | 44.01435 | -91.6158 |
| MUR002 | SW | 43.99247 | -95.9896 |

**Table S2. List of compounds discovered in 2025.** SMILES and InChiKeys are included to identify chemical compounds in common metabolomic databases.

| **Compound** | **Smiles** | **Inchikey** |
| --- | --- | --- |
| 4-Hydroxy-beta-thujone | CC(C)[C@@]12C[C@H]1[C@@](C(=O)C2)(C)O | QCIUNUUHBWFYTD-SFGNSQDASA-N |
| 7-Methoxycoumarin | COC1=CC2=C(C=C1)C=CC(=O)O2 | LIIALPBMIOVAHH-UHFFFAOYSA-N |
| alpha-Copaene | CC1=CC[C@H]2[C@H]3C1[C@]2(CC[C@H]3C(C)C)C | VLXDPFLIRFYIME-XIQJJJERSA-N |
| alpha-Longipinene | CC1=CC[C@@H]2[C@@H]3[C@H]1[C@]2(CCCC3(C)C)C | HICYDYJTCDBHMZ-COMQUAJESA-N |
| alpha-Phellandrene | CC1=CCC(C=C1)C(C)C | OGLDWXZKYODSOB-UHFFFAOYSA-N |
| alpha-Pinene | CC1=CCC2CC1C2(C)C | GRWFGVWFFZKLTI-UHFFFAOYSA-N |
| alpha-Terpineol | CC1=CCC(CC1)C(C)(C)O | WUOACPNHFRMFPN-UHFFFAOYSA-N |
| alpha-Thujenal | CC1=CC(C2(C1C2)C(C)C)C=O | GXVIUCBGLYAPLV-UHFFFAOYSA-N |
| alpha-Thujene | CC1=CCC2(C1C2)C(C)C | KQAZVFVOEIRWHN-UHFFFAOYSA-N |
| alpha-Thujone | C[C@@H]1[C@H]2C[C@]2(CC1=O)C(C)C | USMNOWBWPHYOEA-MRTMQBJTSA-N |
| beta-Bourbonene | CC(C)[C@@H]1CC[C@@]2([C@H]1[C@@H]3[C@H]2CCC3=C)C | YIRAHEODBQONHI-ZQNQSHIBSA-N |
| beta-Copaene | CC(C)[C@@H]1CC[C@]2([C@@H]3[C@H]1C2C(=C)CC3)C | UPVZPMJSRSWJHQ-XIQJJJERSA-N |
| beta-Thujone | C[C@H]1[C@H]2C[C@]2(CC1=O)C(C)C | USMNOWBWPHYOEA-XKSSXDPKSA-N |
| Bicyclogermacrene | C/C/1=C\\CC/C(=C/[C@H]2[C@H](C2(C)C)CC1)/C | VPDZRSSKICPUEY-JEPMYXAXSA-N |
| Bicyclosesquiphellandrene | C[C@@H]1CC[C@H](C2=CC(=C)CC[C@H]12)C(C)C | RNDFUOKDULDZPR-HZSPNIEDSA-N |
| Borneol acetate | CC(=O)O[C@@H]1C[C@@H]2CC[C@]1(C2(C)C)C | KGEKLUUHTZCSIP-HOSYDEDBSA-N |
| Camphene | CC1(C2CCC(C2)C1=C)C | CRPUJAZIXJMDBK-UHFFFAOYSA-N |
| Car-3-en-5-one | CC1=CC(=O)C2C(C1)C2(C)C | WDILKLCBAXJFIA-UHFFFAOYSA-N |
| Caryophyllene | C/C/1=C\\CCC(=C)[C@H]2CC([C@@H]2CC1)(C)C | NPNUFJAVOOONJE-GFUGXAQUSA-N |
| Chrysanthenone | CC1=CCC2C(=O)C1C2(C)C | IECBDTGWSQNQID-UHFFFAOYSA-N |
| cis-Chrysanthenol | CC1=CCC2C(C1C2(C)C)O | IRZWAJHUWGZMMT-UHFFFAOYSA-N |
| cis-Chrysanthenol acetate | CC1=CCC2C(C1C2(C)C)OC(=O)C | UASZOTVHPVEMQR-UHFFFAOYSA-N |
| cis-Cubenol | C[C@H]1CC[C@H]([C@H]2[C@@]1(CCC(=C2)C)O)C(C)C | COGPRPSWSKLKTF-AJNGGQMLSA-N |
| cis-Sabinol | CC(C)[C@@]12C[C@@H]1C(=C)[C@@H](C2)O | MDFQXBNVOAKNAY-BBBLOLIVSA-N |
| Coumarin | C1=CC=C2C(=C1)C=CC(=O)O2 | ZYGHJZDHTFUPRJ-UHFFFAOYSA-N |
| D-Limonene | CC1=CC[C@@H](CC1)C(=C)C | XMGQYMWWDOXHJM-JTQLQIEISA-N |
| Eucalyptol | CC1(C2CCC(O1)(CC2)C)C | WEEGYLXZBRQIMU-UHFFFAOYSA-N |
| Eugenol | COC1=C(C=CC(=C1)CC=C)O | RRAFCDWBNXTKKO-UHFFFAOYSA-N |
| gamma-Cadinene | CC1=C[C@@H]2[C@@H](CC1)C(=C)CC[C@H]2C(C)C | WRHGORWNJGOVQY-KKUMJFAQSA-N |
| gamma-Muurolene | CC1=C[C@@H]2[C@H](CC1)C(=C)CC[C@H]2C(C)C | WRHGORWNJGOVQY-ZNMIVQPWSA-N |
| gamma-Terpinene | CC1=CCC(=CC1)C(C)C | YKFLAYDHMOASIY-UHFFFAOYSA-N |
| Germacrene D | C/C/1=C\\CCC(=C)/C=C/[C@@H](CC1)C(C)C | GAIBLDCXCZKKJE-RXJOXMPGSA-N |
| Isogermacrene D | C/C/1=C\\CCC(=C)/C=C/[C@@H](CC1)C(C)C | GAIBLDCXCZKKJE-RXJOXMPGSA-N |
| Longiverbenone | CC1=CC(=O)C2C3C1C2(CCCC3(C)C)C | KTPOZFYJWLGJGH-UHFFFAOYSA-N |
| Neophytadiene (Isomer #1) | CC(C)CCCC(C)CCCC(C)CC/C=C(\\C)/C=C | HNTNJYMWJHGCBD-LDADJPATSA-N |
| Neophytadiene (Isomer #2) | CC(C)CCCC(C)CCCC(C)CC/C=C(\\C)/C=C | HNTNJYMWJHGCBD-LDADJPATSA-N |
| p-Cymene | CC1=CC=C(C=C1)C(C)C | HFPZCAJZSCWRBC-UHFFFAOYSA-N |
| Pinocarvone | CC1(C2CC1C(=C)C(=O)C2)C | TZDMGBLPGZXHJI-UHFFFAOYSA-N |
| Sabinene | CC(C)C12CCC(=C)C1C2 | NDVASEGYNIMXJL-UHFFFAOYSA-N |
| Sabinene hydrate | CC(C)C12CCC(C1C2)(C)O | KXSDPILWMGFJMM-UHFFFAOYSA-N |
| Spirojatamol | CC(C)[C@@H]1CCC(=C)[C@@]2(C1)CCC[C@]2(C)O | AEUBOEQPIBOTGP-ILXRZTDVSA-N |
| trans-Sabinene hydrate | CC(C)[C@@]12CC[C@@]([C@@H]1C2)(C)O | KXSDPILWMGFJMM-AEJSXWLSSA-N |
| endo-Borneol | C[C@@]12CC[C@@H](C1(C)C)C[C@@H]2O | DTGKSKDOIYIVQL-WEDXCCLWSA-N |
| beta-Pinene | CC1(C2CCC(=C)C1C2)C | WTARULDDTDQWMU-UHFFFAOYSA-N |
| Caryophyllene.oxide | C[C@@]12CC[C@@H]3[C@H](CC3(C)C)C(=C)CC[C@H]1O2 | NVEQFIOZRFFVFW-RGCMKSIDSA-N |
| Germacrene.D.4.ol | C/C/1=C\\CCC(/C=C/C(CC1)C(C)C)(C)O | HCTXHCNRLCYBN-BMCYRRRCSA-N |
| Isophytol | CC(C)CCCC(C)CCCC(C)CCCC(C)(C=C)O | KEVYVLWNCKMXJX-UHFFFAOYSA-N |
| Phytol | C[C@@H](CCC[C@@H](C)CCC/C(=C/CO)/C)CCCC(C)C | BOTWFXYSPFMFNR-PYDDKJGSSA-N |
| Tomentosin | C[C@H]1C[C@@H]2[C@H](CC=C1CCC(=O)C)C(=C)C(=O)O2 | AVFIYMSJDDGDBQ-CUOATXAZSA-N |
| gamma.Elemene | CC(=C1CC[C@]([C@H](C1)C(=C)C)(C)C=C)C | BQSLMQNYHVFRDT-CABCVRRESA-N |
| Trimethyl.3.methylene.hexadeca.1.6.10.14.tetraene | CC(=CCC/C(=C/CC/C(=C/CCC(=C)C=C)/C)/C)C | SIVJVJUIXOEPW-YTOAGJIJSA-N |

**Table S3. Terpene profile PERMANOVA results.** PERMANOVA of compounds using a Euclidean distance matrix evaluated with “Type II” sum of squares. Significant P-values are bolded and denoted as follows: +*P* < 0.1, **P* < 0.05, ***P* < 0.01, ****P* < 0.001.

|  | **Sum Sq** | **Mean Sq** | **Df** | **F** | **P-value** |
| --- | --- | --- | --- | --- | --- |
| Transplant site | 2897.3 | 321.92 | 3 | 7.986 | **0.001**** |
| Source population | 32.8 | 32.81 | 9 | 0.814 | 0.582 |
| Treatment | 34.8 | 34.76 | 1 | 0.862 | 0.53 |
| Transplant site × source population | 74.5 | 24.82 | 27 | 0.616 | 0.944 |
| Transplant site × treatment | 358.8 | 39.87 | 3 | 0.989 | 0.473 |
| Source population × treatment | 242.4 | 60.60 | 9 | 1.503 | **0.05*** |
| Transplant site × block | 231.9 | 57.98 | 4 | 1.438 | *0.066+* |
| Transplant site × source population × treatment | 996.3 | 36.90 | 27 | 0.915 | 0.815 |
| Residual | 9190.6 | 40.31 | 228 | NA | NA |
| Total | 15861 | 51 | 311 | NA | NA |

**Table S4. Pairwise comparison of terpene profiles in climate change treatments.** Pairwise PERMANOVA comparison of compounds using a Euclidean distance matrix evaluated with “Type II” sum of squares.

| **Pairs** | **Sum Sq** | **F** | **R2** | **P-value** | **P-adj** | **sig** |
| --- | --- | --- | --- | --- | --- | --- |
| Heat_STL037 vs Heat_MUR002 | 63.15 | 1.15 | 0.04 | 0.30 | 0.31 |  |
| Heat_STL037 vs Heat_WAB006 | 64.66 | 1.21 | 0.04 | 0.29 | 0.30 |  |
| Heat_STL037 vs Heat_LINC004 | 123.18 | 2.17 | 0.07 | 0.09 | 0.10 |  |
| Heat_STL037 vs Heat_COOK001.5 | 92.35 | 1.78 | 0.05 | 0.09 | 0.10 |  |
| Heat_STL037 vs Heat_PEN002 | 180.83 | 3.10 | 0.09 | 0.00 | 0.01 | * |
| Heat_STL037 vs Amb_PEN002 | 196.18 | 3.34 | 0.10 | 0.00 | 0.00 | * |
| Heat_STL037 vs Amb_BIG002 | 228.44 | 3.41 | 0.11 | 0.03 | 0.04 | . |
| Heat_STL037 vs Amb_LINC004 | 81.38 | 1.43 | 0.04 | 0.19 | 0.21 |  |
| Heat_STL037 vs Amb_STL037 | 46.38 | 0.86 | 0.03 | 0.51 | 0.52 |  |
| Heat_STL037 vs Amb_WIN020 | 310.56 | 5.11 | 0.15 | 0.01 | 0.01 | * |
| Heat_STL037 vs Heat_MARS001 | 159.58 | 2.84 | 0.09 | 0.01 | 0.02 | . |
| Heat_STL037 vs Heat_BIG002 | 179.10 | 3.37 | 0.10 | 0.01 | 0.01 | * |
| Heat_STL037 vs Heat_WIN020 | 294.43 | 5.05 | 0.14 | 0.00 | 0.00 | * |
| Heat_STL037 vs Amb_COOK001.5 | 72.16 | 1.32 | 0.04 | 0.21 | 0.22 |  |
| Heat_STL037 vs Amb_STL035 | 133.98 | 2.25 | 0.07 | 0.07 | 0.08 |  |
| Heat_STL037 vs Amb_MARS001 | 185.72 | 3.36 | 0.10 | 0.00 | 0.01 | * |
| Heat_STL037 vs Amb_WAB006 | 81.98 | 1.61 | 0.05 | 0.15 | 0.16 |  |
| Heat_STL037 vs Amb_MUR002 | 125.35 | 2.57 | 0.08 | 0.02 | 0.03 | . |
| Heat_STL037 vs Heat_STL035 | 111.47 | 1.94 | 0.06 | 0.08 | 0.10 |  |
| Heat_MUR002 vs Heat_WAB006 | 34.75 | 0.92 | 0.03 | 0.49 | 0.50 |  |
| Heat_MUR002 vs Heat_LINC004 | 62.14 | 1.51 | 0.05 | 0.14 | 0.16 |  |
| Heat_MUR002 vs Heat_COOK001.5 | 63.00 | 1.74 | 0.05 | 0.07 | 0.09 |  |
| Heat_MUR002 vs Heat_PEN002 | 218.39 | 5.11 | 0.15 | 0.00 | 0.00 | * |
| Heat_MUR002 vs Amb_PEN002 | 171.77 | 3.97 | 0.12 | 0.00 | 0.00 | * |
| Heat_MUR002 vs Amb_BIG002 | 217.57 | 4.29 | 0.13 | 0.00 | 0.01 | * |
| Heat_MUR002 vs Amb_LINC004 | 56.41 | 1.36 | 0.04 | 0.20 | 0.22 |  |
| Heat_MUR002 vs Amb_STL037 | 53.65 | 1.40 | 0.04 | 0.15 | 0.17 |  |
| Heat_MUR002 vs Amb_WIN020 | 187.72 | 4.19 | 0.13 | 0.01 | 0.01 | * |
| Heat_MUR002 vs Heat_MARS001 | 168.83 | 4.21 | 0.13 | 0.00 | 0.00 | * |
| Heat_MUR002 vs Heat_BIG002 | 145.56 | 3.89 | 0.11 | 0.00 | 0.00 | * |
| Heat_MUR002 vs Heat_WIN020 | 153.75 | 3.64 | 0.11 | 0.01 | 0.01 | . |
| Heat_MUR002 vs Amb_COOK001.5 | 52.04 | 1.33 | 0.04 | 0.20 | 0.22 |  |
| Heat_MUR002 vs Amb_STL035 | 117.68 | 2.70 | 0.09 | 0.02 | 0.03 | . |
| Heat_MUR002 vs Amb_MARS001 | 182.62 | 4.68 | 0.14 | 0.00 | 0.00 | * |
| Heat_MUR002 vs Amb_WAB006 | 60.38 | 1.72 | 0.05 | 0.07 | 0.09 |  |
| Heat_MUR002 vs Amb_MUR002 | 41.05 | 1.24 | 0.04 | 0.27 | 0.29 |  |
| Heat_MUR002 vs Heat_STL035 | 79.38 | 1.94 | 0.06 | 0.06 | 0.08 |  |
| Heat_WAB006 vs Heat_LINC004 | 87.44 | 2.19 | 0.07 | 0.05 | 0.07 |  |
| Heat_WAB006 vs Heat_COOK001.5 | 92.33 | 2.63 | 0.08 | 0.01 | 0.01 | . |
| Heat_WAB006 vs Heat_PEN002 | 210.84 | 5.07 | 0.14 | 0.00 | 0.00 | * |
| Heat_WAB006 vs Amb_PEN002 | 195.32 | 4.64 | 0.13 | 0.00 | 0.00 | * |
| Heat_WAB006 vs Amb_BIG002 | 161.35 | 3.26 | 0.10 | 0.01 | 0.02 | . |
| Heat_WAB006 vs Amb_LINC004 | 53.80 | 1.34 | 0.04 | 0.22 | 0.23 |  |
| Heat_WAB006 vs Amb_STL037 | 65.35 | 1.76 | 0.06 | 0.07 | 0.09 |  |
| Heat_WAB006 vs Amb_WIN020 | 248.34 | 5.69 | 0.16 | 0.00 | 0.01 | * |
| Heat_WAB006 vs Heat_MARS001 | 193.87 | 4.99 | 0.15 | 0.00 | 0.00 | * |
| Heat_WAB006 vs Heat_BIG002 | 104.61 | 2.89 | 0.09 | 0.01 | 0.02 | . |
| Heat_WAB006 vs Heat_WIN020 | 221.33 | 5.40 | 0.16 | 0.00 | 0.00 | * |
| Heat_WAB006 vs Amb_COOK001.5 | 66.86 | 1.76 | 0.06 | 0.07 | 0.09 |  |
| Heat_WAB006 vs Amb_STL035 | 163.72 | 3.86 | 0.12 | 0.01 | 0.01 | . |
| Heat_WAB006 vs Amb_MARS001 | 225.55 | 5.96 | 0.17 | 0.00 | 0.00 | * |
| Heat_WAB006 vs Amb_WAB006 | 35.81 | 1.05 | 0.03 | 0.38 | 0.40 |  |
| Heat_WAB006 vs Amb_MUR002 | 63.20 | 1.99 | 0.06 | 0.05 | 0.06 |  |
| Heat_WAB006 vs Heat_STL035 | 111.60 | 2.81 | 0.09 | 0.01 | 0.02 | . |
| Heat_LINC004 vs Heat_COOK001.5 | 81.55 | 2.13 | 0.07 | 0.06 | 0.07 |  |
| Heat_LINC004 vs Heat_PEN002 | 298.72 | 6.67 | 0.18 | 0.00 | 0.00 | * |
| Heat_LINC004 vs Amb_PEN002 | 235.98 | 5.21 | 0.15 | 0.00 | 0.00 | * |
| Heat_LINC004 vs Amb_BIG002 | 284.83 | 5.39 | 0.16 | 0.00 | 0.00 | * |
| Heat_LINC004 vs Amb_LINC004 | 75.47 | 1.74 | 0.05 | 0.09 | 0.10 |  |
| Heat_LINC004 vs Amb_STL037 | 98.34 | 2.44 | 0.08 | 0.04 | 0.05 | . |
| Heat_LINC004 vs Amb_WIN020 | 142.46 | 3.04 | 0.09 | 0.02 | 0.03 | . |
| Heat_LINC004 vs Heat_MARS001 | 160.85 | 3.82 | 0.12 | 0.00 | 0.01 | * |
| Heat_LINC004 vs Heat_BIG002 | 189.07 | 4.80 | 0.14 | 0.00 | 0.00 | * |
| Heat_LINC004 vs Heat_WIN020 | 122.26 | 2.76 | 0.09 | 0.03 | 0.04 | . |
| Heat_LINC004 vs Amb_COOK001.5 | 87.57 | 2.13 | 0.07 | 0.03 | 0.04 | . |
| Heat_LINC004 vs Amb_STL035 | 105.12 | 2.30 | 0.07 | 0.04 | 0.05 |  |
| Heat_LINC004 vs Amb_MARS001 | 160.98 | 3.92 | 0.12 | 0.00 | 0.01 | * |
| Heat_LINC004 vs Amb_WAB006 | 141.47 | 3.80 | 0.11 | 0.01 | 0.01 | * |
| Heat_LINC004 vs Amb_MUR002 | 82.84 | 2.37 | 0.07 | 0.03 | 0.04 | . |
| Heat_LINC004 vs Heat_STL035 | 58.18 | 1.35 | 0.05 | 0.22 | 0.24 |  |
| Heat_COOK001.5 vs Heat_PEN002 | 302.86 | 7.58 | 0.20 | 0.00 | 0.00 | * |
| Heat_COOK001.5 vs Amb_PEN002 | 250.88 | 6.20 | 0.17 | 0.00 | 0.00 | * |
| Heat_COOK001.5 vs Amb_BIG002 | 264.53 | 5.54 | 0.17 | 0.00 | 0.00 | * |
| Heat_COOK001.5 vs Amb_LINC004 | 102.38 | 2.65 | 0.08 | 0.01 | 0.01 | * |
| Heat_COOK001.5 vs Amb_STL037 | 63.79 | 1.79 | 0.06 | 0.07 | 0.08 |  |
| Heat_COOK001.5 vs Amb_WIN020 | 298.87 | 7.13 | 0.20 | 0.00 | 0.00 | * |
| Heat_COOK001.5 vs Heat_MARS001 | 222.23 | 5.98 | 0.17 | 0.00 | 0.00 | * |
| Heat_COOK001.5 vs Heat_BIG002 | 191.69 | 5.54 | 0.16 | 0.00 | 0.00 | * |
| Heat_COOK001.5 vs Heat_WIN020 | 222.27 | 5.65 | 0.16 | 0.00 | 0.00 | * |
| Heat_COOK001.5 vs Amb_COOK001.5 | 21.73 | 0.60 | 0.02 | 0.85 | 0.85 |  |
| Heat_COOK001.5 vs Amb_STL035 | 189.98 | 4.67 | 0.14 | 0.00 | 0.00 | * |
| Heat_COOK001.5 vs Amb_MARS001 | 236.90 | 6.56 | 0.18 | 0.00 | 0.00 | * |
| Heat_COOK001.5 vs Amb_WAB006 | 84.31 | 2.61 | 0.08 | 0.01 | 0.02 | . |
| Heat_COOK001.5 vs Amb_MUR002 | 95.65 | 3.17 | 0.10 | 0.01 | 0.01 | * |
| Heat_COOK001.5 vs Heat_STL035 | 138.14 | 3.64 | 0.12 | 0.00 | 0.00 | * |
| Heat_PEN002 vs Amb_PEN002 | 42.59 | 0.91 | 0.03 | 0.53 | 0.55 |  |
| Heat_PEN002 vs Amb_BIG002 | 322.12 | 5.89 | 0.17 | 0.00 | 0.00 | * |
| Heat_PEN002 vs Amb_LINC004 | 120.40 | 2.67 | 0.08 | 0.01 | 0.01 | . |
| Heat_PEN002 vs Amb_STL037 | 202.68 | 4.82 | 0.14 | 0.00 | 0.00 | * |
| Heat_PEN002 vs Amb_WIN020 | 365.28 | 7.51 | 0.21 | 0.00 | 0.00 | * |
| Heat_PEN002 vs Heat_MARS001 | 184.93 | 4.21 | 0.13 | 0.00 | 0.00 | * |
| Heat_PEN002 vs Heat_BIG002 | 293.65 | 7.15 | 0.19 | 0.00 | 0.00 | * |
| Heat_PEN002 vs Heat_WIN020 | 409.51 | 8.90 | 0.23 | 0.00 | 0.00 | * |
| Heat_PEN002 vs Amb_COOK001.5 | 271.92 | 6.36 | 0.17 | 0.00 | 0.00 | * |
| Heat_PEN002 vs Amb_STL035 | 199.45 | 4.21 | 0.13 | 0.00 | 0.00 | * |
| Heat_PEN002 vs Amb_MARS001 | 288.09 | 6.73 | 0.19 | 0.00 | 0.00 | * |
| Heat_PEN002 vs Amb_WAB006 | 256.71 | 6.61 | 0.18 | 0.00 | 0.00 | * |
| Heat_PEN002 vs Amb_MUR002 | 347.22 | 9.47 | 0.24 | 0.00 | 0.00 | * |
| Heat_PEN002 vs Heat_STL035 | 214.22 | 4.77 | 0.15 | 0.00 | 0.00 | * |
| Amb_PEN002 vs Amb_BIG002 | 315.63 | 5.71 | 0.17 | 0.00 | 0.00 | * |
| Amb_PEN002 vs Amb_LINC004 | 98.58 | 2.16 | 0.07 | 0.02 | 0.03 | . |
| Amb_PEN002 vs Amb_STL037 | 161.75 | 3.80 | 0.11 | 0.00 | 0.00 | * |
| Amb_PEN002 vs Amb_WIN020 | 278.08 | 5.65 | 0.16 | 0.00 | 0.00 | * |
| Amb_PEN002 vs Heat_MARS001 | 175.71 | 3.96 | 0.12 | 0.00 | 0.01 | * |
| Amb_PEN002 vs Heat_BIG002 | 268.27 | 6.45 | 0.18 | 0.00 | 0.00 | * |
| Amb_PEN002 vs Heat_WIN020 | 297.53 | 6.39 | 0.18 | 0.00 | 0.00 | * |
| Amb_PEN002 vs Amb_COOK001.5 | 217.25 | 5.02 | 0.14 | 0.00 | 0.00 | * |
| Amb_PEN002 vs Amb_STL035 | 194.86 | 4.06 | 0.12 | 0.00 | 0.00 | * |
| Amb_PEN002 vs Amb_MARS001 | 257.20 | 5.93 | 0.17 | 0.00 | 0.00 | * |
| Amb_PEN002 vs Amb_WAB006 | 223.36 | 5.67 | 0.16 | 0.00 | 0.00 | * |
| Amb_PEN002 vs Amb_MUR002 | 261.06 | 7.02 | 0.19 | 0.00 | 0.00 | * |
| Amb_PEN002 vs Heat_STL035 | 176.71 | 3.89 | 0.12 | 0.00 | 0.00 | * |
| Amb_BIG002 vs Amb_LINC004 | 183.80 | 3.45 | 0.11 | 0.01 | 0.01 | * |
| Amb_BIG002 vs Amb_STL037 | 206.36 | 4.13 | 0.13 | 0.00 | 0.00 | * |
| Amb_BIG002 vs Amb_WIN020 | 448.57 | 7.82 | 0.22 | 0.00 | 0.00 | * |
| Amb_BIG002 vs Heat_MARS001 | 323.32 | 6.19 | 0.19 | 0.00 | 0.00 | * |
| Amb_BIG002 vs Heat_BIG002 | 25.08 | 0.51 | 0.02 | 0.82 | 0.82 |  |
| Amb_BIG002 vs Heat_WIN020 | 414.99 | 7.61 | 0.22 | 0.00 | 0.00 | * |
| Amb_BIG002 vs Amb_COOK001.5 | 211.84 | 4.18 | 0.13 | 0.00 | 0.00 | * |
| Amb_BIG002 vs Amb_STL035 | 365.63 | 6.53 | 0.19 | 0.00 | 0.00 | * |
| Amb_BIG002 vs Amb_MARS001 | 380.57 | 7.45 | 0.22 | 0.00 | 0.00 | * |
| Amb_BIG002 vs Amb_WAB006 | 178.36 | 3.83 | 0.12 | 0.00 | 0.01 | * |
| Amb_BIG002 vs Amb_MUR002 | 227.04 | 5.14 | 0.16 | 0.00 | 0.00 | * |
| Amb_BIG002 vs Heat_STL035 | 290.40 | 5.42 | 0.17 | 0.00 | 0.00 | * |
| Amb_LINC004 vs Amb_STL037 | 81.75 | 2.01 | 0.06 | 0.05 | 0.06 |  |
| Amb_LINC004 vs Amb_WIN020 | 188.79 | 3.99 | 0.12 | 0.01 | 0.01 | * |
| Amb_LINC004 vs Heat_MARS001 | 122.45 | 2.88 | 0.09 | 0.01 | 0.01 | * |
| Amb_LINC004 vs Heat_BIG002 | 111.91 | 2.81 | 0.09 | 0.01 | 0.02 | . |
| Amb_LINC004 vs Heat_WIN020 | 179.59 | 4.02 | 0.12 | 0.00 | 0.00 | * |
| Amb_LINC004 vs Amb_COOK001.5 | 83.82 | 2.02 | 0.06 | 0.03 | 0.04 | . |
| Amb_LINC004 vs Amb_STL035 | 100.03 | 2.17 | 0.07 | 0.05 | 0.06 |  |
| Amb_LINC004 vs Amb_MARS001 | 161.62 | 3.90 | 0.12 | 0.00 | 0.00 | * |
| Amb_LINC004 vs Amb_WAB006 | 108.17 | 2.88 | 0.09 | 0.01 | 0.01 | . |
| Amb_LINC004 vs Amb_MUR002 | 122.19 | 3.46 | 0.10 | 0.00 | 0.00 | * |
| Amb_LINC004 vs Heat_STL035 | 78.79 | 1.81 | 0.06 | 0.08 | 0.09 |  |
| Amb_STL037 vs Amb_WIN020 | 251.62 | 5.71 | 0.16 | 0.00 | 0.00 | * |
| Amb_STL037 vs Heat_MARS001 | 182.40 | 4.64 | 0.14 | 0.00 | 0.00 | * |
| Amb_STL037 vs Heat_BIG002 | 145.67 | 3.97 | 0.12 | 0.00 | 0.00 | * |
| Amb_STL037 vs Heat_WIN020 | 207.74 | 5.01 | 0.15 | 0.00 | 0.01 | * |
| Amb_STL037 vs Amb_COOK001.5 | 30.11 | 0.79 | 0.03 | 0.66 | 0.67 |  |
| Amb_STL037 vs Amb_STL035 | 168.05 | 3.92 | 0.12 | 0.01 | 0.01 | . |
| Amb_STL037 vs Amb_MARS001 | 208.70 | 5.45 | 0.16 | 0.00 | 0.00 | * |
| Amb_STL037 vs Amb_WAB006 | 52.28 | 1.52 | 0.05 | 0.12 | 0.14 |  |
| Amb_STL037 vs Amb_MUR002 | 70.43 | 2.18 | 0.07 | 0.02 | 0.03 | . |
| Amb_STL037 vs Heat_STL035 | 102.52 | 2.55 | 0.08 | 0.02 | 0.03 | . |
| Amb_WIN020 vs Heat_MARS001 | 243.35 | 5.28 | 0.16 | 0.00 | 0.00 | * |
| Amb_WIN020 vs Heat_BIG002 | 340.34 | 7.90 | 0.21 | 0.00 | 0.00 | * |
| Amb_WIN020 vs Heat_WIN020 | 43.18 | 0.89 | 0.03 | 0.45 | 0.47 |  |
| Amb_WIN020 vs Amb_COOK001.5 | 290.72 | 6.49 | 0.18 | 0.00 | 0.00 | * |
| Amb_WIN020 vs Amb_STL035 | 108.53 | 2.18 | 0.07 | 0.07 | 0.09 |  |
| Amb_WIN020 vs Amb_MARS001 | 234.51 | 5.21 | 0.16 | 0.00 | 0.00 | * |
| Amb_WIN020 vs Amb_WAB006 | 349.88 | 8.57 | 0.23 | 0.00 | 0.00 | * |
| Amb_WIN020 vs Amb_MUR002 | 216.64 | 5.62 | 0.16 | 0.00 | 0.00 | * |
| Amb_WIN020 vs Heat_STL035 | 77.47 | 1.64 | 0.06 | 0.12 | 0.14 |  |
| Heat_MARS001 vs Heat_BIG002 | 266.32 | 6.95 | 0.19 | 0.00 | 0.00 | * |
| Heat_MARS001 vs Heat_WIN020 | 273.90 | 6.32 | 0.18 | 0.00 | 0.00 | * |
| Heat_MARS001 vs Amb_COOK001.5 | 211.92 | 5.29 | 0.15 | 0.00 | 0.00 | * |
| Heat_MARS001 vs Amb_STL035 | 110.82 | 2.47 | 0.08 | 0.02 | 0.03 | . |
| Heat_MARS001 vs Amb_MARS001 | 34.50 | 0.86 | 0.03 | 0.56 | 0.57 |  |
| Heat_MARS001 vs Amb_WAB006 | 274.03 | 7.61 | 0.21 | 0.00 | 0.00 | * |
| Heat_MARS001 vs Amb_MUR002 | 262.12 | 7.77 | 0.21 | 0.00 | 0.00 | * |
| Heat_MARS001 vs Heat_STL035 | 133.36 | 3.17 | 0.11 | 0.00 | 0.00 | * |
| Heat_BIG002 vs Heat_WIN020 | 306.34 | 7.57 | 0.21 | 0.00 | 0.00 | * |
| Heat_BIG002 vs Amb_COOK001.5 | 141.52 | 3.79 | 0.11 | 0.00 | 0.00 | * |
| Heat_BIG002 vs Amb_STL035 | 281.85 | 6.73 | 0.19 | 0.00 | 0.00 | * |
| Heat_BIG002 vs Amb_MARS001 | 317.16 | 8.51 | 0.23 | 0.00 | 0.00 | * |
| Heat_BIG002 vs Amb_WAB006 | 134.53 | 4.02 | 0.12 | 0.00 | 0.00 | * |
| Heat_BIG002 vs Amb_MUR002 | 159.54 | 5.10 | 0.15 | 0.00 | 0.00 | * |
| Heat_BIG002 vs Heat_STL035 | 214.19 | 5.47 | 0.16 | 0.00 | 0.00 | * |
| Heat_WIN020 vs Amb_COOK001.5 | 220.58 | 5.23 | 0.15 | 0.00 | 0.00 | * |
| Heat_WIN020 vs Amb_STL035 | 162.08 | 3.45 | 0.11 | 0.01 | 0.01 | * |
| Heat_WIN020 vs Amb_MARS001 | 253.26 | 5.99 | 0.18 | 0.00 | 0.00 | * |
| Heat_WIN020 vs Amb_WAB006 | 280.02 | 7.33 | 0.20 | 0.00 | 0.00 | * |
| Heat_WIN020 vs Amb_MUR002 | 170.31 | 4.74 | 0.14 | 0.00 | 0.00 | * |
| Heat_WIN020 vs Heat_STL035 | 102.04 | 2.30 | 0.08 | 0.03 | 0.03 | . |
| Amb_COOK001.5 vs Amb_STL035 | 192.78 | 4.42 | 0.13 | 0.01 | 0.01 | * |
| Amb_COOK001.5 vs Amb_MARS001 | 221.29 | 5.67 | 0.16 | 0.00 | 0.00 | * |
| Amb_COOK001.5 vs Amb_WAB006 | 55.03 | 1.57 | 0.05 | 0.11 | 0.13 |  |
| Amb_COOK001.5 vs Amb_MUR002 | 70.61 | 2.14 | 0.07 | 0.03 | 0.04 | . |
| Amb_COOK001.5 vs Heat_STL035 | 126.78 | 3.10 | 0.10 | 0.01 | 0.01 | . |
| Amb_STL035 vs Amb_MARS001 | 119.39 | 2.73 | 0.09 | 0.01 | 0.01 | . |
| Amb_STL035 vs Amb_WAB006 | 258.44 | 6.53 | 0.18 | 0.00 | 0.00 | * |
| Amb_STL035 vs Amb_MUR002 | 211.95 | 5.68 | 0.16 | 0.00 | 0.00 | * |
| Amb_STL035 vs Heat_STL035 | 31.75 | 0.69 | 0.02 | 0.69 | 0.70 |  |
| Amb_MARS001 vs Amb_WAB006 | 300.89 | 8.60 | 0.23 | 0.00 | 0.00 | * |
| Amb_MARS001 vs Amb_MUR002 | 257.65 | 7.88 | 0.21 | 0.00 | 0.00 | * |
| Amb_MARS001 vs Heat_STL035 | 125.97 | 3.08 | 0.10 | 0.01 | 0.02 | . |
| Amb_WAB006 vs Amb_MUR002 | 79.96 | 2.75 | 0.08 | 0.01 | 0.02 | . |
| Amb_WAB006 vs Heat_STL035 | 176.13 | 4.79 | 0.15 | 0.00 | 0.00 | * |
| Amb_MUR002 vs Heat_STL035 | 121.47 | 3.53 | 0.11 | 0.00 | 0.00 | * |

**Table S5. Confusion matrices to evaluate region within similarities of pairwise population treatment comparisons.** We evaluated patterns of significance from the pairwise PERMANOVA comparison of compounds using confusion matrices. “Match” designation within the table refers to significant pairwise comparisons between populations within a region whereas “not match” indicates populations from different regions.

| **Confusion Matrices** | | | |
| --- | --- | --- | --- |
| ***Treatment*** | | | |
|  | Match | Not Match | Total |
| Significant | 72 | 74 | 146 |
| Non-Significant | 18 | 26 | 44 |
|  | 90 | 100 | 190 |
| ***Region*** | | | |
|  | Match | Not Match | Total |
| Significant | 22 | 124 | 146 |
| Non-Significant | 18 | 26 | 44 |
|  | 40 | 150 | 190 |

**Table S6. Primary dominant compounds found in each chemotype as identified by Z-scores extracted from hierarchical clustering.**

| **Chemotype** | **Dominant Compound** | **Rank** | **Average Z-Score** |
| --- | --- | --- | --- |
| 1 | cis.Chrysanthenol.acetate | 1 | 0.704 |
| 1 | endo.Borneol | 2 | 0.589 |
| 1 | alpha.Thujene | 3 | 0.587 |
| 1 | cis.Chrysanthenol | 4 | 0.549 |
| 1 | Borneol.acetate | 5 | 0.538 |
| 2 | beta.Thujone | 1 | 0.702 |
| 2 | X4.Hydroxy.beta.thujone | 2 | 0.676 |
| 2 | alpha.Thujone | 3 | 0.615 |
| 2 | Longiverbenone | 4 | 0.435 |
| 2 | Germacrene.D | 5 | 0.433 |
| 3 | alpha.Thujone | 1 | 0.365 |
| 3 | beta.Thujone | 2 | 0.302 |
| 3 | X4.Hydroxy.beta.thujone | 3 | 0.148 |
| 3 | X7.Methoxycoumarin | 4 | -0.105 |
| 3 | Coumarin | 5 | -0.195 |
| 4 | Car.3.en.5.one | 1 | 2.168 |
| 4 | Chrysanthenone | 2 | 2.158 |
| 4 | alpha.Thujenal | 3 | 1.892 |
| 4 | alpha.Pinene | 4 | 1.321 |
| 4 | Coumarin | 5 | 1.191 |
| 5 | beta.Pinene | 1 | 1.338 |
| 5 | alpha.Thujene | 2 | 1.225 |
| 5 | Camphene | 3 | 1.105 |
| 5 | alpha.Pinene | 4 | 1.078 |
| 5 | Pinocarvone | 5 | 1.072 |

**Table S7. Top 15 dominant compounds found in each chemotypes as identified from important features of the PLS-DA.** Accuracy = 0.77, Q2 = 0.29161, R2 = 0.33585.

|  |  |  | Chemotype | | | | |
| --- | --- | --- | --- | --- | --- | --- | --- |
|  | Compounds Class | Mean Coefficient | 1 | 2 | 3 | 4 | 5 |
| alpha.Thujenal | MTs | 34.4 | 27.2 | 29.8 | 14.8 | 80.3 | 19.9 |
| alpha.Pinene | MTs | 32.7 | 22.9 | 29.5 | 22.6 | 57.2 | 31.4 |
| Chrysanthenone | MTs | 32.2 | 25.1 | 20.1 | 10.3 | 92.2 | 13.6 |
| Car.3.en.5.one | MTs | 30.5 | 21.6 | 9.3 | 9.3 | 100.0 | 12.2 |
| endo.Borneol | MTs | 30.2 | 22.9 | 33.2 | 20.3 | 46.4 | 28.3 |
| X4.Hydroxy.beta.thujone | MTs | 29.5 | 26.7 | 44.3 | 15.6 | 39.1 | 21.6 |
| beta.Thujone | MTs | 28.9 | 25.0 | 45.4 | 20.0 | 26.2 | 28.1 |
| alpha.Thujone | MTs | 27.2 | 24.2 | 40.5 | 15.2 | 34.8 | 21.1 |
| alpha.Thujene | MTs | 27.0 | 17.8 | 27.6 | 23.3 | 33.3 | 32.9 |
| Camphene | MTs | 26.9 | 20.0 | 33.5 | 21.6 | 29.1 | 30.4 |
| beta.Pinene | MTs | 26.2 | 15.7 | 23.2 | 24.8 | 32.1 | 35.1 |
| cis.Chrysanthenol.acetate | MTs | 23.4 | 18.2 | 26.9 | 15.0 | 35.7 | 21.0 |
| alpha.Terpineol | MTs | 23.2 | 15.2 | 24.7 | 21.0 | 25.6 | 29.6 |
| Borneol.acetate | MTs | 22.6 | 16.9 | 22.0 | 13.8 | 41.5 | 19.1 |
| trans.Sabinene.hydrate | MTs | 21.6 | 12.7 | 13.8 | 18.2 | 37.6 | 25.7 |
| Sabinene | MTs | 21.2 | 11.9 | 14.8 | 20.0 | 30.8 | 28.4 |
| beta.Bourbonene | SQTs | 21.0 | 19.1 | 37.0 | 13.8 | 15.5 | 19.4 |
| X7.Methoxycoumarin | Coumarins | 20.4 | 16.2 | 14.8 | 6.9 | 54.9 | 9.3 |
| Sabinene.hydrate | MTs | 20.0 | 10.0 | 10.4 | 19.9 | 31.5 | 28.2 |
| Coumarin | Coumarins | 19.7 | 15.7 | 11.9 | 5.1 | 59.2 | 6.7 |
| alpha.Copaene | SQTs | 19.2 | 17.7 | 36.3 | 13.5 | 9.6 | 18.9 |
| Bicyclogermacrene | SQTs | 19.1 | 18.3 | 38.3 | 13.0 | 7.5 | 18.2 |
| Isogermacrene.D | MTs | 18.8 | 14.1 | 18.9 | 11.9 | 32.2 | 16.7 |
| gamma.Muurolene | SQTs | 18.8 | 16.9 | 34.6 | 13.8 | 9.1 | 19.4 |
| Pinocarvone | MTs | 18.6 | 11.1 | 20.1 | 20.0 | 13.7 | 28.4 |
| beta.Copaene | SQTs | 18.2 | 17.1 | 36.9 | 13.7 | 4.2 | 19.2 |
| gamma.Elemene | SQTs | 18.0 | 15.7 | 32.8 | 14.4 | 7.0 | 20.3 |
| gamma.Cadinene | SQTs | 18.0 | 17.5 | 38.4 | 13.2 | 2.4 | 18.6 |
| Germacrene.D | SQTs | 17.8 | 17.5 | 37.4 | 11.7 | 5.8 | 16.3 |
| Isophytol | DTs | 17.2 | 17.9 | 35.9 | 8.8 | 11.4 | 12.1 |
| Neophytadiene..Isomer..1. | MTs | 15.7 | 16.3 | 32.9 | 8.2 | 9.7 | 11.4 |
| cis.Chrysanthenol | DTs | 15.7 | 13.2 | 23.6 | 11.2 | 14.6 | 15.9 |
| Bicyclosesquiphellandrene | SQTs | 15.4 | 14.4 | 32.0 | 12.2 | 1.1 | 17.3 |
| Neophytadiene..Isomer..2. | DTs | 15.4 | 16.2 | 33.3 | 8.0 | 8.2 | 11.2 |
| Eugenol | Phenylpropanoids | 15.2 | 9.0 | 15.4 | 15.7 | 13.5 | 22.4 |
| gamma.Terpinene | MTs | 14.6 | 6.2 | 12.2 | 20.2 | 5.5 | 29.0 |
| alpha.Phellandrene | SQTs | 14.6 | 4.2 | 3.9 | 20.6 | 14.5 | 29.7 |
| D.Limonene | SQTs | 13.4 | 7.7 | 13.8 | 14.9 | 9.4 | 21.4 |
| Caryophyllene | MTs | 13.4 | 15.4 | 34.1 | 6.7 | 1.4 | 9.4 |
| Germacrene.D.4.ol | SQTs | 12.8 | 12.2 | 25.7 | 8.9 | 5.0 | 12.5 |
| Caryophyllene.oxide | SQTs | 12.3 | 7.6 | 7.4 | 8.7 | 25.7 | 12.3 |
| Phytol | DTs | 12.2 | 12.4 | 26.1 | 7.3 | 4.6 | 10.4 |
| alpha.Longipinene | SQTs | 12.1 | 13.7 | 26.1 | 4.2 | 10.5 | 5.9 |
| cis.Sabinol | MTs | 11.9 | 8.4 | 6.4 | 5.8 | 30.4 | 8.3 |
| Longiverbenone | SQTs | 11.8 | 14.8 | 32.0 | 4.1 | 2.0 | 5.8 |
| p.Cymene | MTs | 11.2 | 5.8 | 9.6 | 12.6 | 9.8 | 18.0 |
| Eucalyptol | MTs | 10.5 | 6.3 | 14.3 | 13.0 | 0.0 | 18.8 |
| X.E.E..7.11.15.Trimethyl.3 | Fatty acyls | 8.5 | 8.1 | 16.5 | 5.5 | 4.9 | 7.8 |
| cis.Cubenol | SQTs | 7.5 | 3.4 | 6.8 | 9.8 | 3.2 | 14.2 |
| Tomentosin | SQTs | 5.6 | 2.9 | 3.9 | 5.7 | 6.9 | 8.4 |
| Spirojatamol | SQTs | 4.4 | 1.1 | 1.8 | 6.5 | 3.0 | 9.4 |

**Table S8. Chemotype ANOVA table.** Significant P-values are bolded and denoted as follows: +*P* < 0.1, **P* < 0.05, ***P* < 0.01, ****P* < 0.001.

|  | **Chi-sq** | **Df** | **P-value** |
| --- | --- | --- | --- |
| Transplant site | 10.8 | 3 | **0.013**** |
| Source population | 54.7 | 9 | **0.000***** |
| Treatment | 0.2 | 1 | 0.649 |
| Transplant site × source population | 31.8 | 27 | 0.240 |
| Transplant site × treatment | 0.5 | 3 | 0.929 |
| Source population × treatment | 5.1 | 9 | 0.822 |
| Transplant site × block | 19.4 | 16 | 0.251 |
| Transplant site × source population × treatment | 30.7 | 27 | 0.283 |

**Table S9. PERMANOVA of terpene profiles between years.** PERMANOVA of compounds using a Euclidean distance matrix evaluated with “Type II” sum of squares. Significant P-values are bolded and denoted as follows: +*P* < 0.1, **P* < 0.05, ***P* < 0.01, ****P* < 0.001.

|  | Sum Sq | Mean Sq | Df | F | P-value |
| --- | --- | --- | --- | --- | --- |
| Transplant site | 1796.025 | 199.558 | 1 | 6.045 | **0.001**** |
| Source population | 54.612 | 54.612 | 9 | 1.654 | *0.06+* |
| Year | 54.809 | 54.809 | 1 | 1.66 | *0.073+* |
| Transplant site × source population | 99.491 | 99.491 | 9 | 3.014 | **0.004**** |
| Transplant site × year | 319.661 | 35.518 | 1 | 1.076 | 0.271 |
| Source population × year | 383.14 | 47.892 | 9 | 1.451 | **0.007**** |
| Transplant site × block | 387.863 | 48.483 | 8 | 1.469 | **0.007**** |
| Residual | 4522.532 | 33.011 | 137 | NA | NA |
| Total | 7729.365 | 44.168 | 175 | NA | NA |

**Table S10. ANOVA table for likelihood of herbivory.** The chemotype × source population interaction was not included as not all chemotypes were represented in each source populations, leading to reduced ability to address biological questions, and to reduce model complexity. Significant P-values are bolded and denoted as follows: +*P* < 0.1, **P* < 0.05, ***P* < 0.01, ****P* < 0.001.

|  | **LR Chisq** | **Df** | **P-value** |
| --- | --- | --- | --- |
| Chemotype | 5.25 | 4 | 0.263 |
| Transplant site | 61.84 | 3 | **0.000***** |
| Source population | 8.23 | 9 | 0.511 |
| Treatment | 1.60 | 1 | 0.205 |
| Chemotype × transplant site | 19.03 | 12 | *0.088+* |
| Chemotype × source population | - | - | - |
| Chemotype × treatment | 14.72 | 4 | **0.005**** |
| Transplant site × source population | 28.90 | 27 | 0.366 |
| Transplant site × treatment | 11.78 | 3 | **0.008**** |
| Source population × treatment | 23.36 | 9 | **0.005**** |

**Table S11. ANOVA for chemotype models with latitude.** We centered latitude and longitude (mean = 0) to facilitate comparison of regression coefficients. Significant P-values are bolded and denoted as follows: +*P* < 0.1, **P* < 0.05, ***P* < 0.01, ****P* < 0.001.

|  | **Chisq** | **Df** | **P-value** |
| --- | --- | --- | --- |
| Chemotype | 4.13 | 4.00 | 0.388 |
| Transplant site | 89.98 | 3.00 | **2.21E-19***** |
| Treatment | 2.64 | 1.00 | 0.104 |
| Latitude | 18.27 | 1.00 | **0.000019***** |
| Longitude | 0.31 | 1.00 | 0.577 |
| Chemotype × transplant site | 9.19 | 12.00 | 0.686 |
| Chemotype × treatment | 5.18 | 4.00 | 0.269 |
| Chemotype × latitude | 7.66 | 4.00 | 0.105 |
| Chemotype × longitude | 4.47 | 4.00 | 0.346 |
| Transplant site × treatment | 3.35 | 3.00 | 0.340 |
| Transplant site × latitude | 7.99 | 3.00 | **0.046*** |
| Transplant site × longitude | 6.60 | 3.00 | *0.086+* |
| Treatment × latitude | 0.40 | 1.00 | 0.528 |
| Treatment × longitude | 0.40 | 1.00 | 0.526 |
| Latitude × longitude | 0.45 | 1.00 | 0.504 |

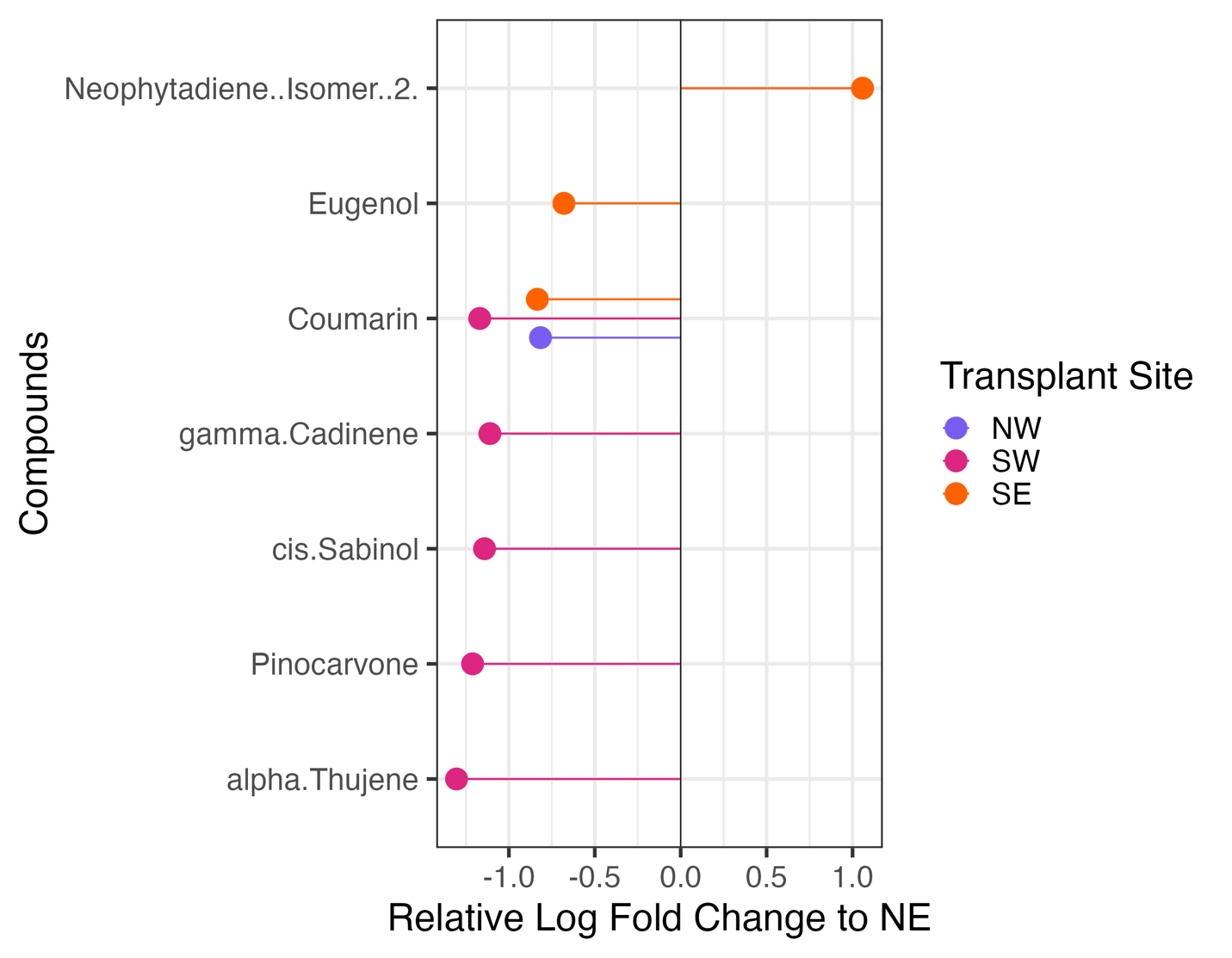

**Fig. S1. Differentially expressed compounds in each transplant site.** Results from ANCOM-BC analysis. Log fold changes are relative to the NE site (negative values = increased expression in the NE site, positive values = increased expression in the comparison site). Colors are transplant sites.

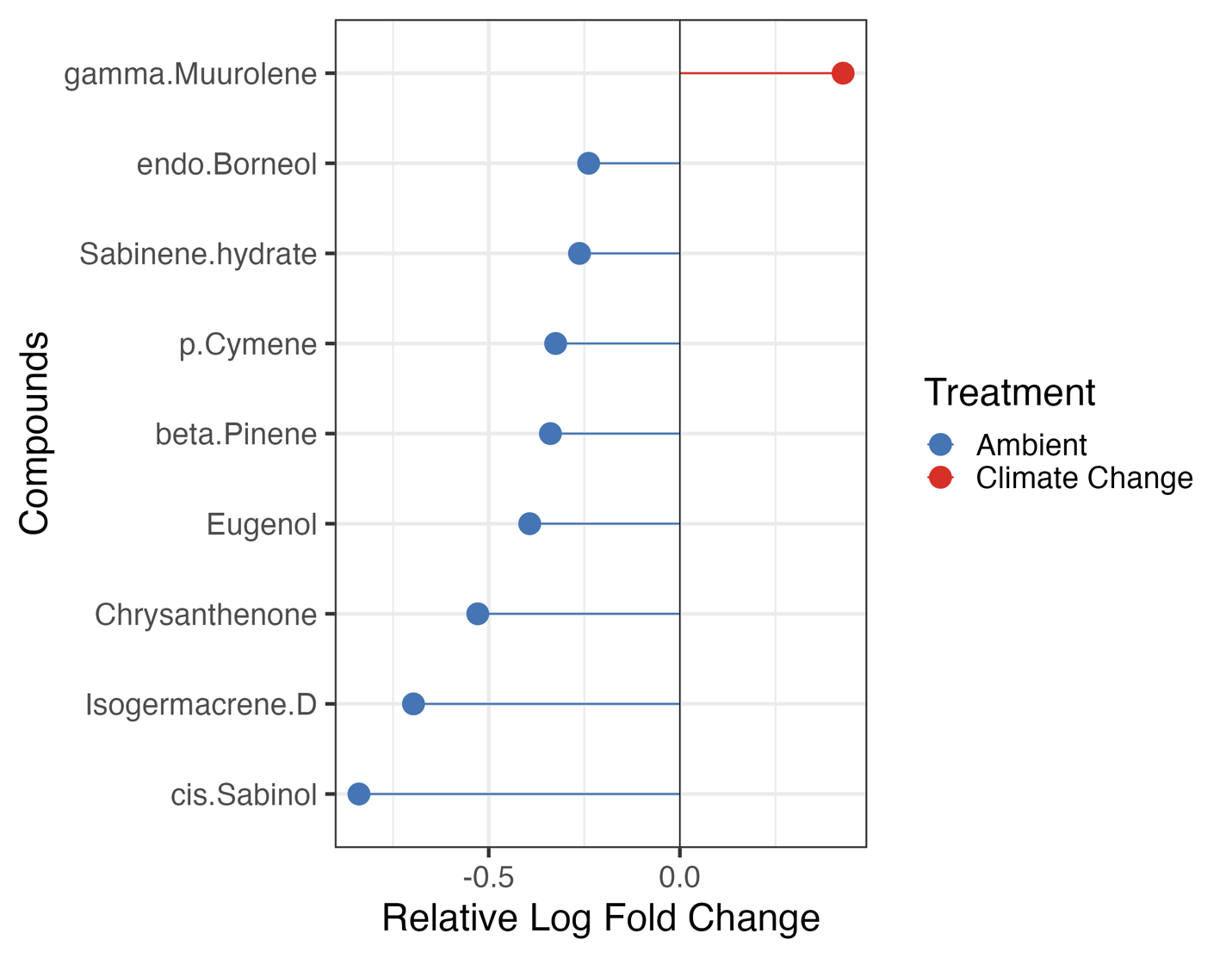

**Fig. S2. Differentially expressed compounds between treatments relative to the ambient treatments.** Log fold changes are relative to the ambient treatment (negative values = increased expression in the ambient treatment, positive values = increased expression in the climate change treatment). Results from ANCOM-BC analysis. Colors are climate change treatments.

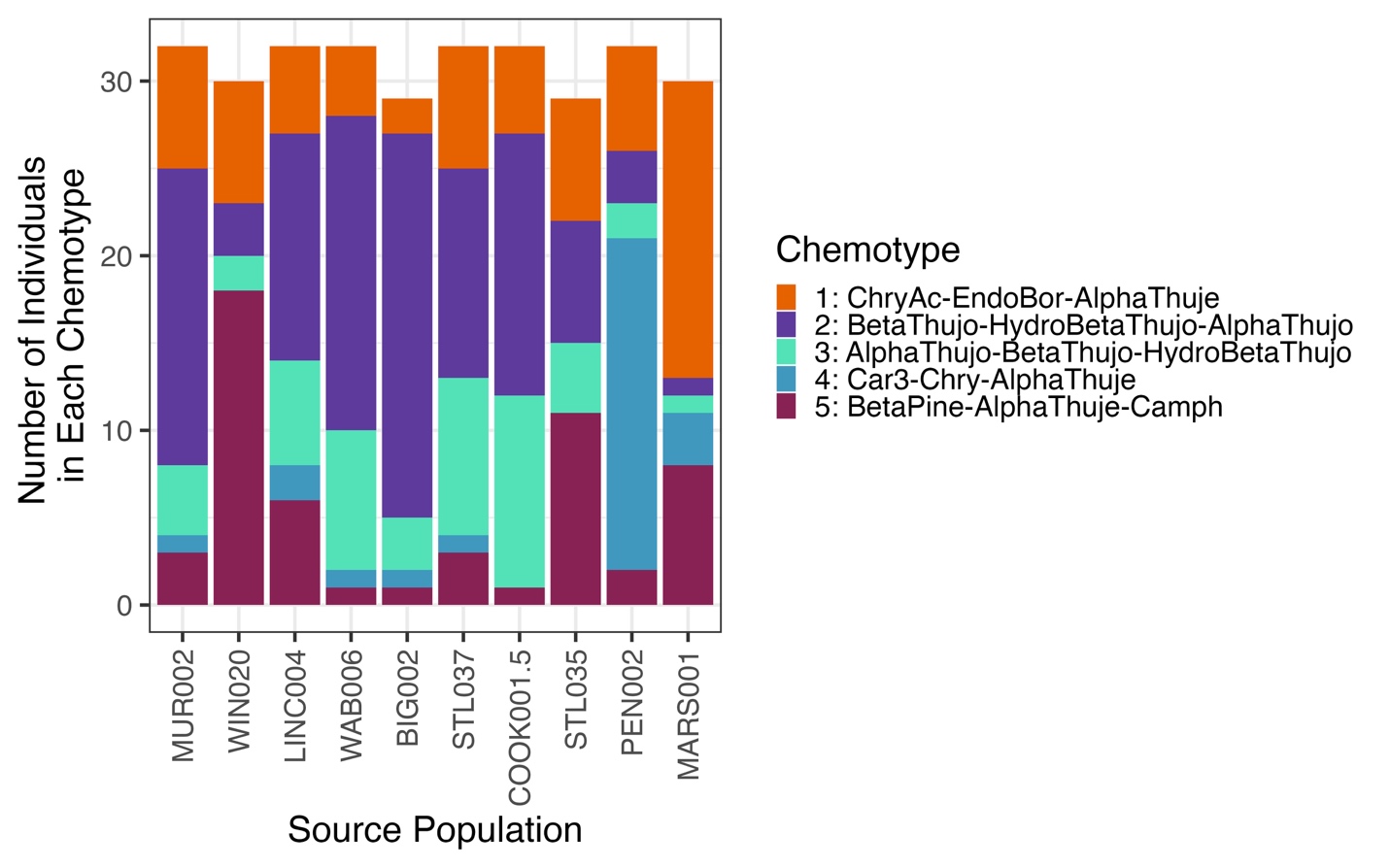

**Fig. S3. Proportion of chemotypes within each population.** Source populations are ordered from S to N. Colors are chemotype groups.

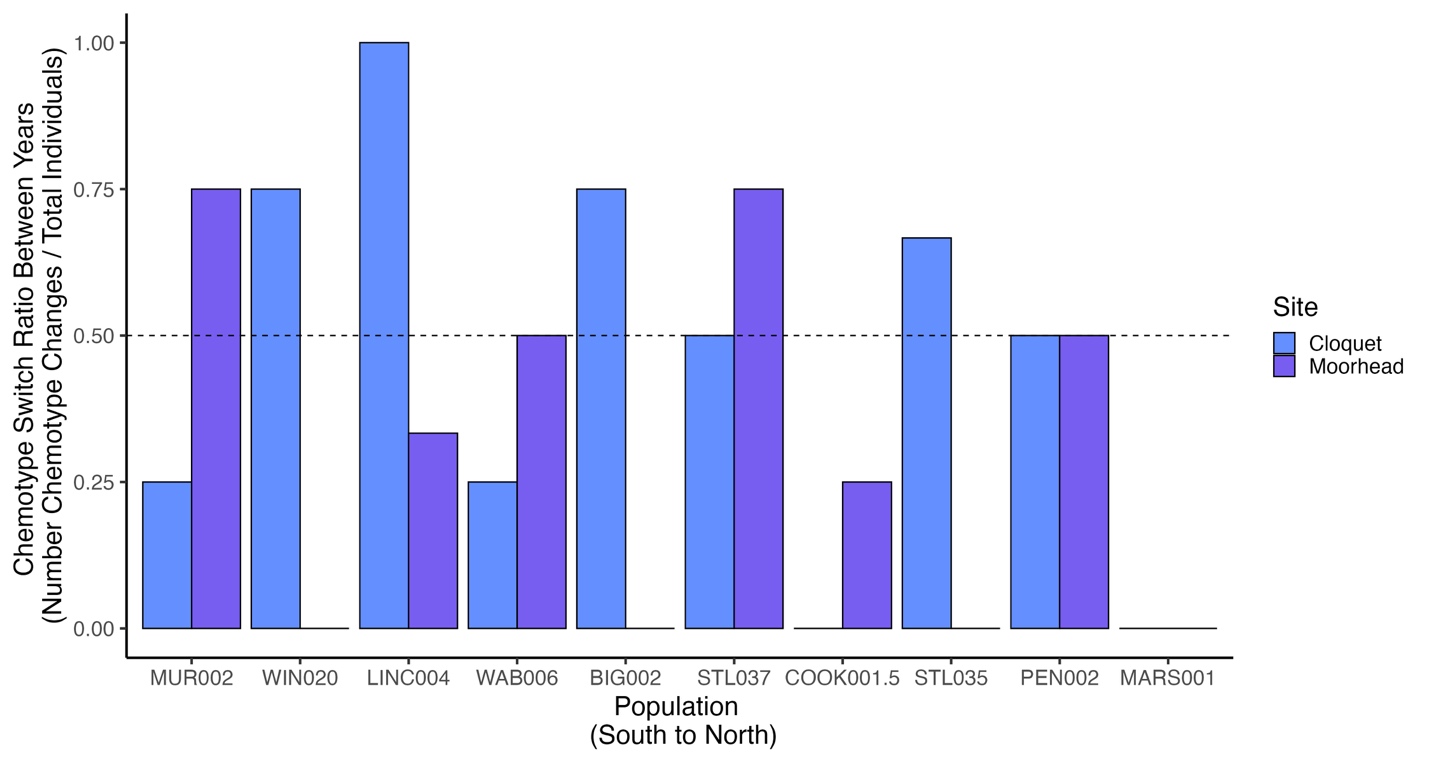

**Fig. S4. Switching chemotypes between years.** Bars represent the number of individuals that switched chemotypes out of the total number of individuals within a given population at either the NE (blue) or NW (purple) transplant sites. Colors are transplant sites. Dashed line at 0.50 is for visual reference.

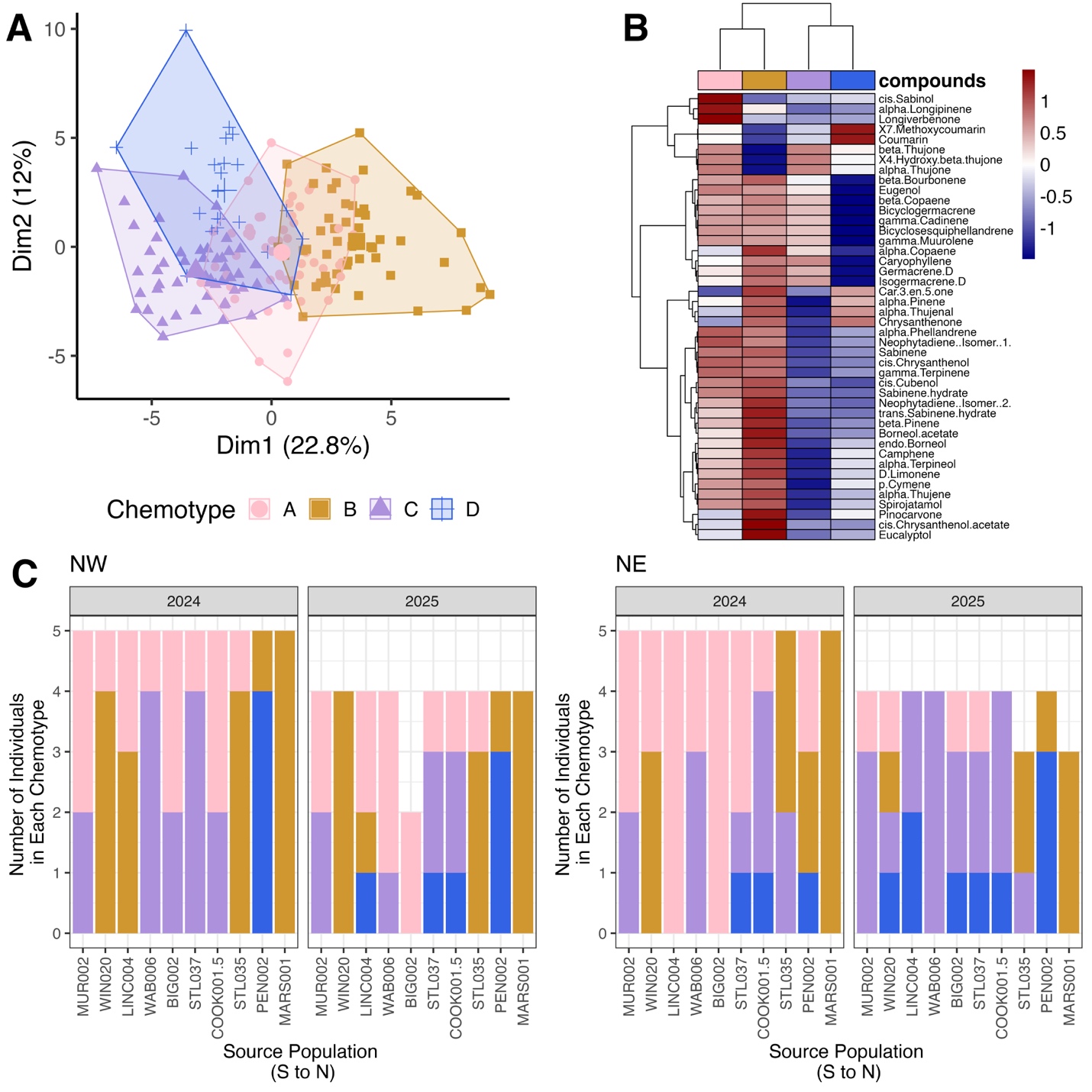

**Fig. S5.** **Chemotypes clustered spatially and temporally in the combined 2024 and 2025 dataset.** These analyses include individuals from the northern transplant sites in the ambient treatment. We denoted these Chemotypes as A, B, C, D to differentiate them from the chemotype groups in the climate change (2025) analyses. Individuals with the unique chemotype (Chemotype A) had a higher relative abundance of cis-sabinol compared to the other chemotypes which primarily determined this grouping. Chemotypes B, C, and D shared two of three dominant compounds with Chemotypes 1, 2, and 4 (respectively) identified in 2025. Points in (A) and counts in (C) are all individuals characterized from 10 populations across four transplant sites. The gradient bar in (B) shows differences in relative abundance of terpenes (red = increased, blue = decreased). Colors are chemotype groups as determined by hierarchical clustering.

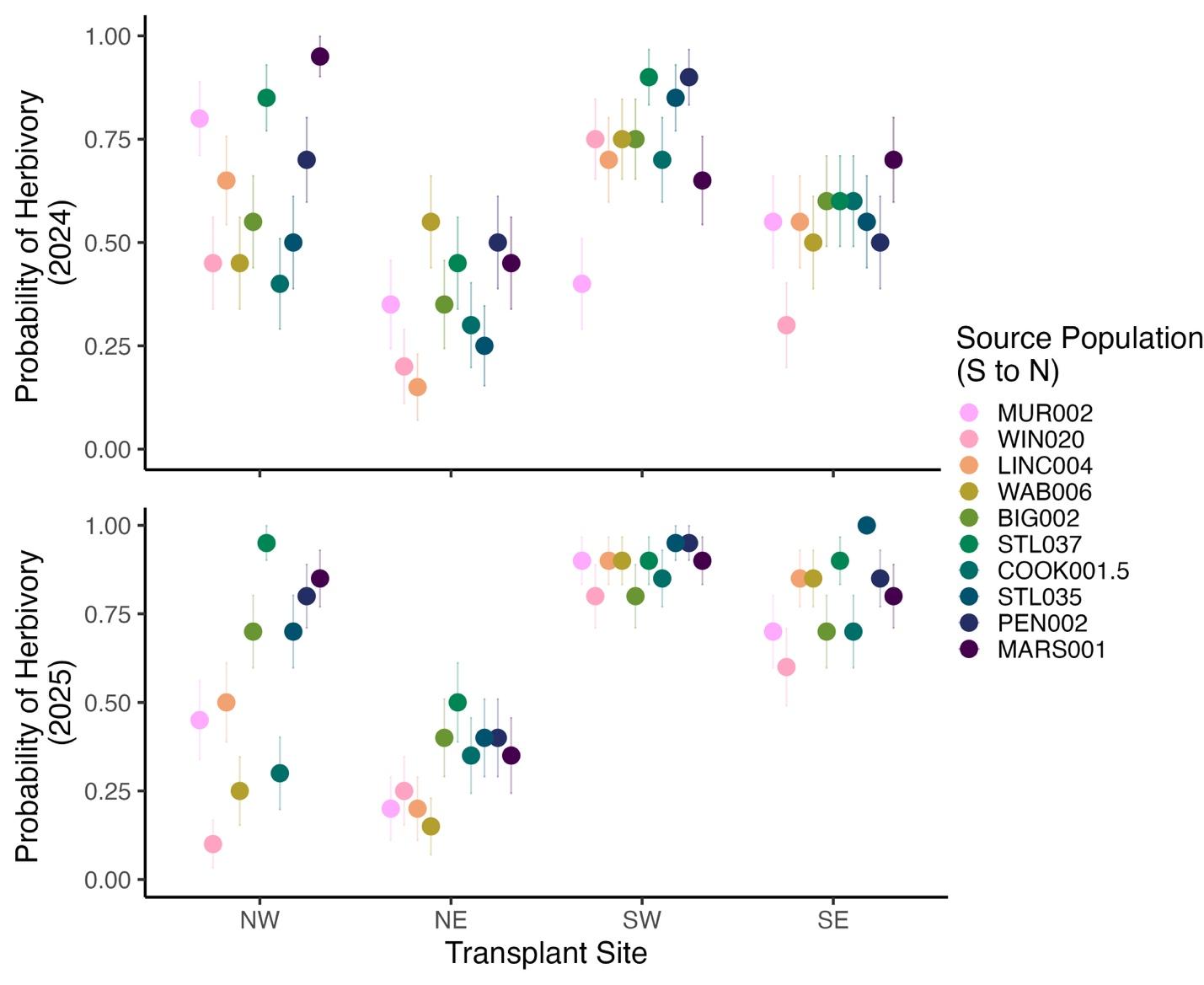

**Fig. S6. The overall probability of herbivory increased at more southern transplant sites, but herbivory differences among populations were apparent at northern transplant sites.** Points are means and bars are standard error. Colors are source populations ordered from south to north.
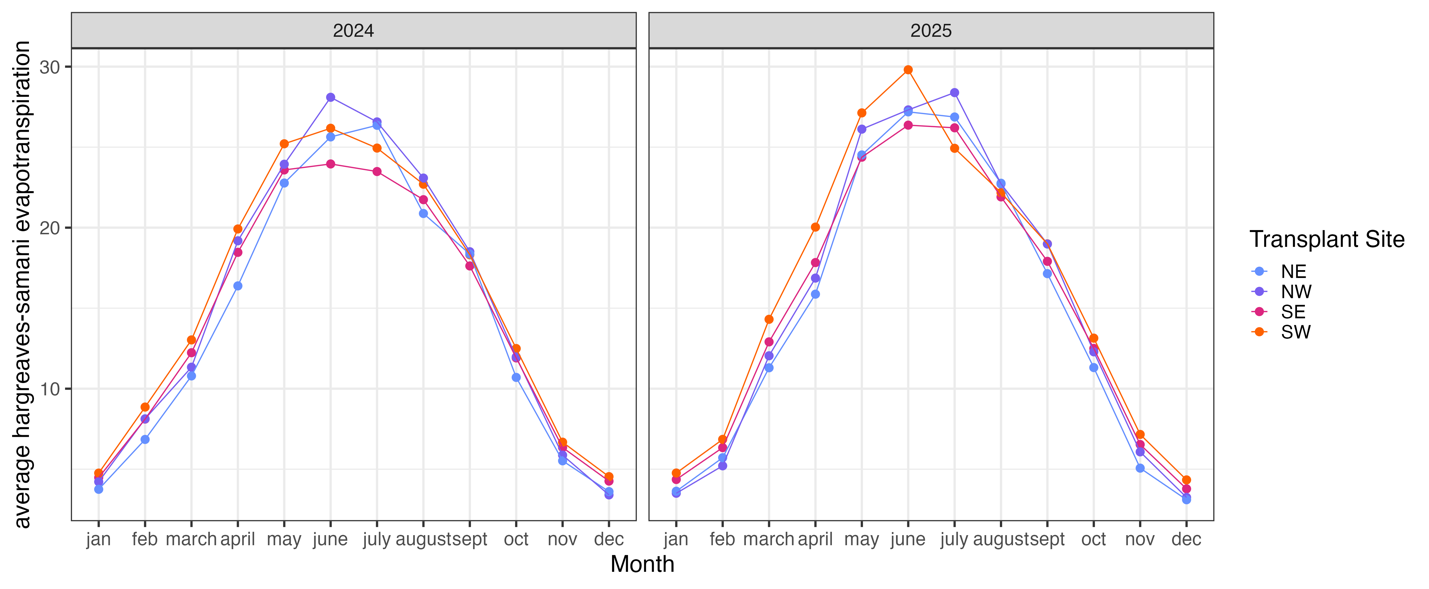

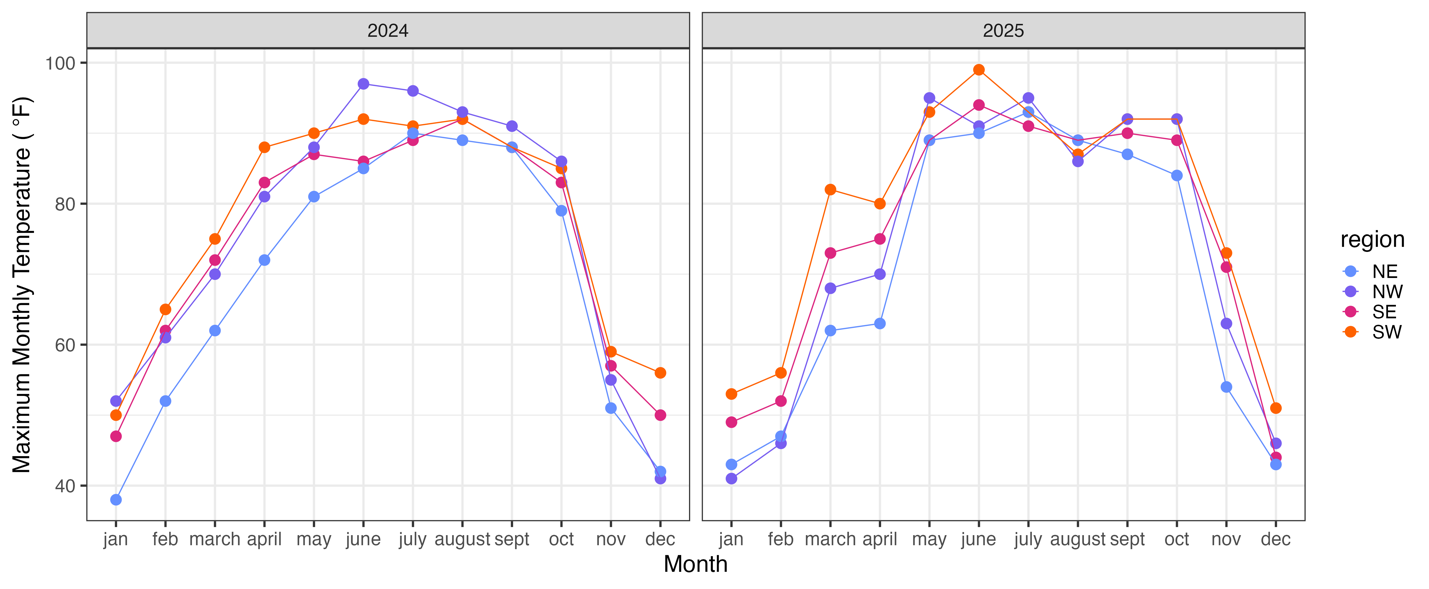

**Fig. S7. Monthly climatic averages during experimental years.** Experiment year climate data were retrieved from on-site (NE, NW, and SE) or nearby (SW) weather stations accessed from the Applied Climate Information System (ACIS) hosted by NOAA National Centers for Environmental Information. Points are monthly averages. Temperature differences are reported in °F and precipitation differences are reported in inches. Colors are transplant site regions.

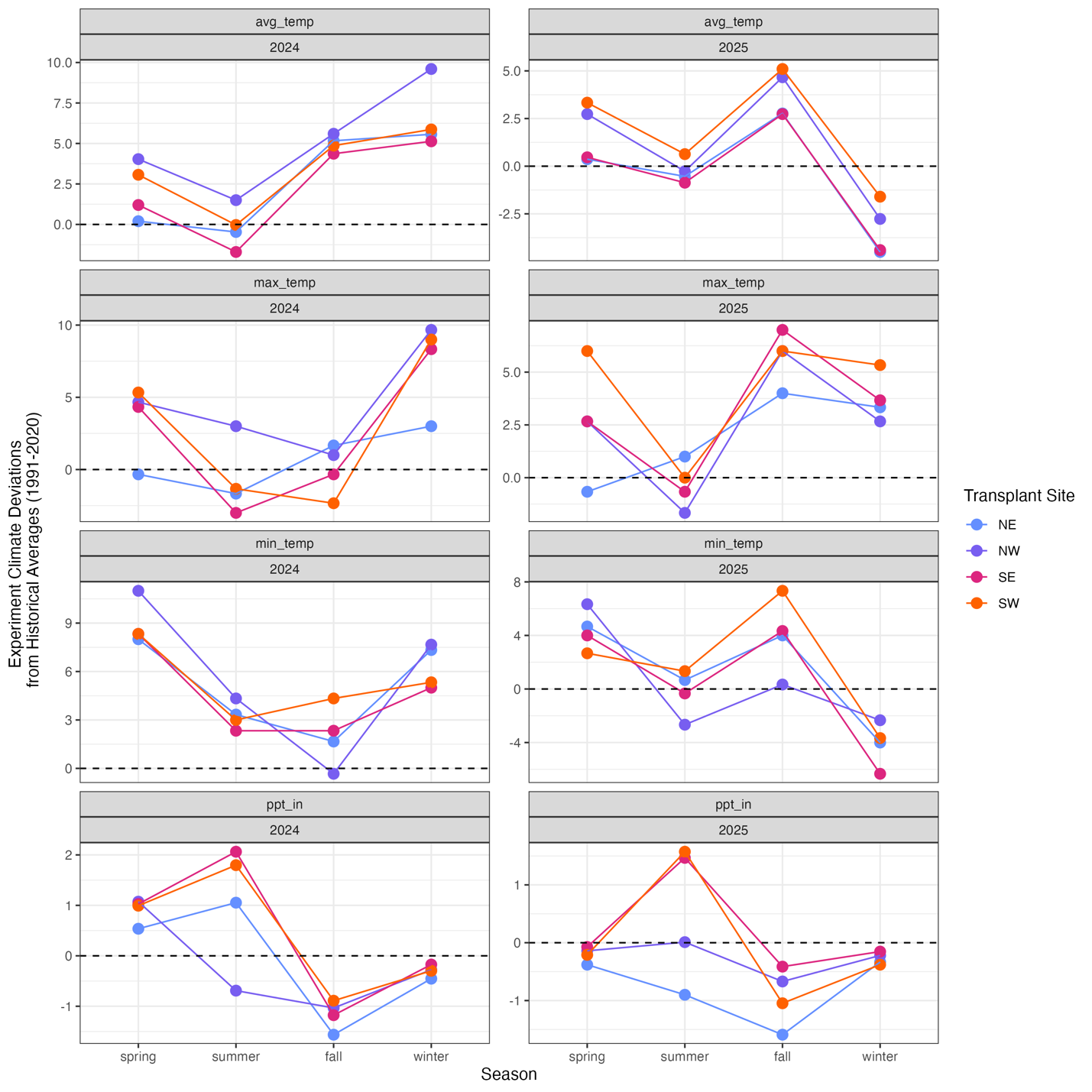

**Fig. S8. Seasonal climate deviations (anomalies) of experiment years (2024, 2025) compared to 30-year historical averages (1990-2020).** Experiment year and historical climate data were retrieved from on-site (NE, NW, and SE) or nearby (SW) weather stations accessed from the Applied Climate Information System (ACIS) hosted by NOAA National Centers for Environmental Information. Points are seasonal averages are seasons are designated as summer (June-August), spring (March-May), winter (Dec-Feb), fall (September-November). Temperature differences are reported in °F and precipitation differences are reported in inches. Colors are transplant site regions. Dashed line is at 0 for visual reference.
